# Meningeal dendritic cells-astrocytes interactions elevate the kynurenine metabolic pathway to sustain neuropathic pain

**DOI:** 10.1101/2021.01.25.428142

**Authors:** Alexandre G. Maganin, Guilherme R. Souza, Miriam D. Fonseca, Alexandre H. Lopes, Rafaela M. Guimarães, André Dagostin, Nerry T. Cecilio, Atlante S. Mendes, Francisco I. Gomes, Lucas M. Marques, Rangel L Silva, Leticia M. Arruda, Denis A. Santana, Henrique Lemos, Lei Huang, Marcela Davoli-Ferreira, Danielle S. Coelho, Morena B. Sant’anna, Ricardo Kusuda, Jhimmy Talbot, Gabriela Pacholczyk, Gabriela A. Buqui, Norberto P. Lopes, Jose C. Alves-Filho, Ricardo Leão, Jason C. O’Connor, Fernando Q. Cunha, Andrew Mellor, Thiago M. Cunha

## Abstract

Neuropathic pain is triggered by injury to the somatosensory system, and is one of the most important types of chronic pain. Nevertheless, critical pathophysiological mechanisms that maintain neuropathic pain are poorly understood. Here, we show that neuropathic pain is abrogated when the kynurenine metabolic pathway (KYNPATH) initiated by the enzyme indoleamine 2,3-dioxygenase (IDO) is ablated pharmacologically or genetically. Mechanistically, it was found that IDO upregulation in dendritic cells that accumulate in the dorsal root leptomeninges led to increased levels of kynurenine (Kyn) in the spinal cord, where Kyn is metabolized by astrocytes-expressed kynurenine-3-monooxygenase into a pro-nociceptive metabolite 3-hydroxykynurenine. In conclusion, these data reveal a novel role for KYNPATH as an important factor maintaining neuropathic pain during neuroimmune-glia cells interactions. This novel paradigm offers potential new targets for drug development against this type of chronic pain.

---

Neuropathic pain is one of the most clinically relevant types of chronic pain. It is generally caused by direct damage to the somatosensory nervous system. Although several pathophysiological mechanisms that generate neuropathic pain have been described, developing effective treatments remain a challenge^1^. Among the mechanisms involved in the development and maintenance of neuropathic pain, neuron-glial cells (microglia/astrocytes/oligodendrocytes) interactions in the spinal cord seem to play a crucial role through the amplification of central sensitization^2^. More specifically, glial cells-derived mediators that enhance glutamatergic transmission in spinal cord neurons are thought to maintain neuropathic pain^2, 3^.

The kynurenine (Kyn) metabolic pathway (KYNPATH) is a catabolic system linked to pathophysiological processes^4, 5^. For instance, disturbances in KYNPATH have been implicated in several human diseases such as depression, schizophrenia, Alzheimer’s and Huntington’s diseases^6, 7^. Bioactive kynurenines (including Kyn itself) are generated by oxidative catabolism of the essential amino acid tryptophan mediated by indoleamine 2,3-dioxygenase (IDO) or tryptophan 2,3-dioxygenase (TDO) in peripheral tissues and the central nervous system (CNS)^8–10^. In addition, Kyn can be converted to several downstream kynurenine metabolites, such as 3-hydroxykynurenine (3-Hk), 3-hydroxyanthranilic acid (3-Haa) and quinolinic acid (QA), which are mainly mediated by two downstream enzymes, kynurenine-3-monooxygenase (KMO) and 3-hydroxyanthranilic acid dioxygenase (HAAO), respectively (Fig. 1a)^11–13^. Kynureninase is also involved in the conversion of 3-Hk into 3-Haa (Fig. 1a)^5^. On the other hand, Kyn can be also metabolized in a side arm by kynurenine aminotransferases (KATs) into kynurenic acid (Kyna) (Fig. 1a)^14^. These “kynurenines” are biologically active in both the periphery and CNS. The upregulation of KYNPATH is generally a consequence of immune and glial cells activation in either the periphery or the CNS, respectively^4, 15^. Elevated levels of kynurenine metabolites in the CNS, especially 3-Hk and QA might cause neuronal disfunction and damage, which are dependent on an increase in oxidative stress and NMDA glutamatergic receptors activation, respectively^16, 17^. Although the participation of KYNPATH in the pathophysiology of neuropathic pain has been investigated, these few studies produced discrepant results^18–20^. Furthermore, none of them deeply investigated the cellular and molecular underlying mechanisms.

**Fig. 1.**
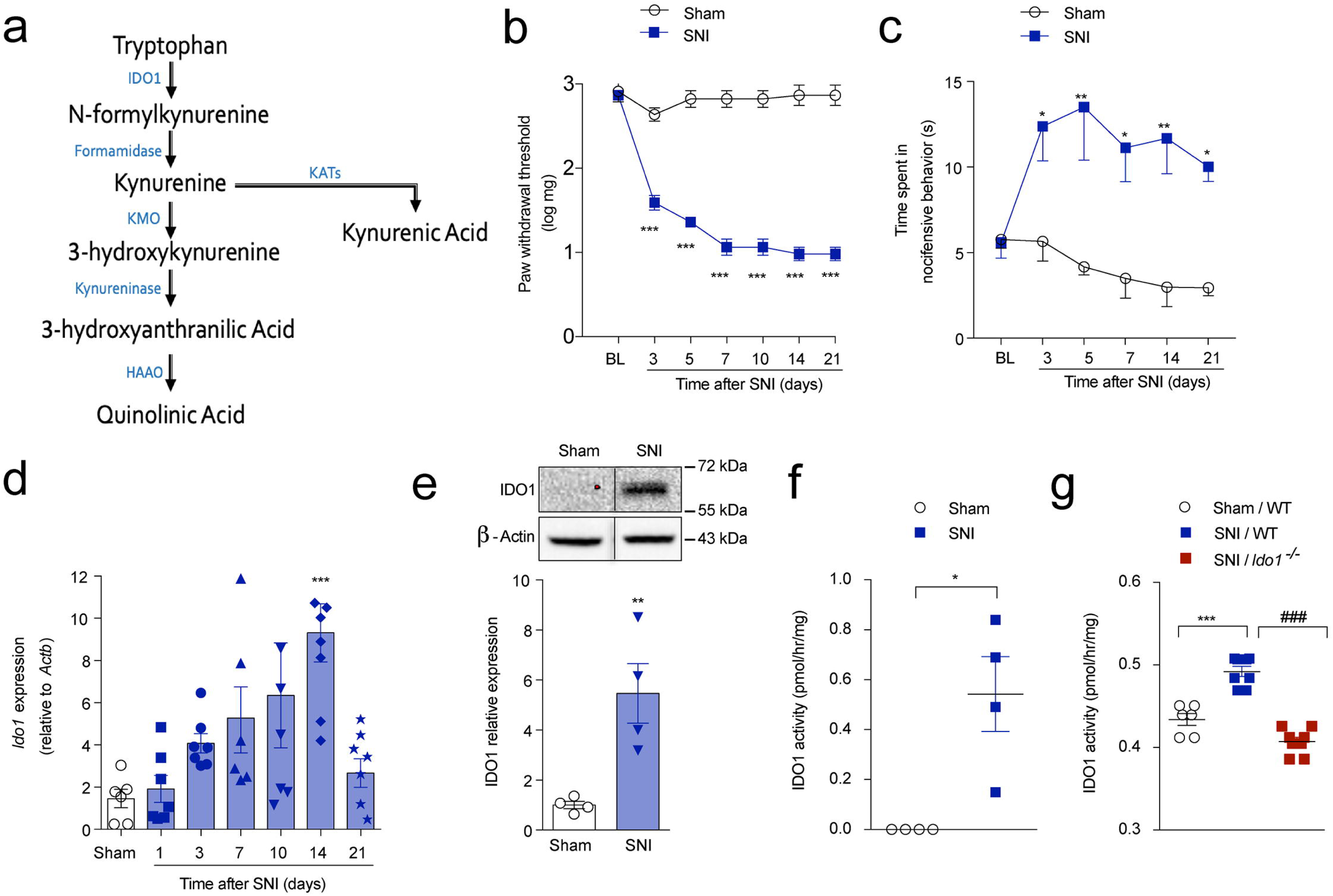
IDO1 expression and activity in spinal cord after peripheral nerve injury. **(a)** Simplified diagram of the kynurenine metabolic pathway. Time-course of **(b)** mechanical or **(c)** cold allodynia after spared nerve injury (SNI) model (n=7). **(d)** *Ido1* mRNA expression in the ipsilateral dorsal horn of the spinal cord after sham (14 days) or SNI surgeries (3-21 day after injury) (n=6-8) **(e)** Western blotting analysis of IDO1 expression in the ipsilateral dorsal horn of the spinal cord 14 days after sham or SNI surgeries (n=5) **(f)** IDO1 enzymatic activity in the ipsilateral dorsal horn of the spinal cords 14 days after sham or SNI surgeries in WT (n=4, pooled 5) and **(g)** compared to *Ido1*^-/-^ mice (n=6-8 pooled of 5). Data are expressed as the mean ± s.e.m. \**P*< 0.05, \*\**P*< 0.01, \*\*\**P*< 0.001 versus sham or ###*P*< 0.001 versus *Ido1^-/-^* mice. Two-way ANOVA, Bonferroni’s post-test (b and c); One-way ANOVA Bonferroni’s post-test (d, and g); Unpaired Student’s *t-*test (e and f).

Considering these points, in the present study we used a series of genetic, biochemical and pharmacological approaches to investigate the role of KYNPATH for the development of neuropathic pain. Here, we show that the neuropathic pain is abrogated when the KYNPATH is pharmacologically or genetically ablated. Mechanistically, peripheral nerve injury induced an upregulation of IDO1 in dendritic cells (DCs) that accumulate into the dorsal root leptomeninges (DRL) leading to an increase in the levels of Kyn in the spinal cord. In the spinal cord, Kyn is metabolized by astrocytes-expressed KMO into a pronociceptive metabolite 3-Hk. These findings elucidate the crucial role of KYNPATH in the maintenance of neuropathic pain and also pointed out novel targets for the development of drugs for neuropathic pain control.

## Results

### Peripheral nerve injury-induced neuropathic pain depends on IDO1

Peripheral nerve injury triggers the activation of glial cells in the spinal cord which participate in the development of neuropathic pain through the production of a vast range of pro-nociceptive mediators^21, 22^. Given that KYNPATH, specially IDO1 is generally up-regulated by inflammatory mediators^23^, we firstly hypothesized that the neuro-immune response in the spinal cord after peripheral nerve injury could promote the upregulation of IDO1 and consequently the KYNPATH that in turn could participate in development of neuropathic pain. To test this initial hypothesis, we used a well-established model of peripheral nerve injury-induced neuropathic pain, the spared nerve injury model (SNI)^24^ which produced mechanical and cold allodynia from 3 days after nerve injury until experimental endpoints (day 21), as well as a robust activation of microglial cells in the spinal cord when compared to sham-operated mice (Fig. 1b and c**, and Supplementary Fig. 1**). The development of SNI-induced neuropathic pain was associated with increased expression (mRNA and protein) and activity of IDO1 in the ipsilateral dorsal horn of spinal cord (Fig. 1d-g). To examine the involvement of IDO1 in neuropathic pain development, firstly we tested the phenotype of *Ido1* null (–/–) mice in SNI-induced mechanical and cold allodynia. Notably, *Ido1*^-/-^ mice developed mechanical allodynia at earlier stage after SNI (3 days after surgery), however, pain hypersensitivity began to ameliorate from day 7 to 28 after surgery (Fig. 2a). However, no difference was observed in SNI-induced cold allodynia between *Ido1*^-/-^ and wild type (WT) mice (Fig. 2b). Importantly, *Ido1*^-/-^ mice exhibited normal thermal pain thresholds (over temperatures ranging from 48-56°C) compared to WT mice (Fig. 2c). Furthermore, compared to WT mice, *Ido1*^-/-^ mice also showed no difference in either total nociceptive behaviour in the first and second phases after the formalin injection (Fig. 2d) or the carrageenan-induced mechanical inflammatory pain hypersensitivity (Fig. 2e). Besides, Complete Freund Adjuvant (CFA)-induced mechanical and thermal pain hypersensitivity in *Ido1* ^-/-^ mice did not differ from to WT mice (Fig 2 f,g). Secondly, WT mice were pharmacologically treated once, at peak of IDO1 expression (14 days after SNI), with two different IDO1 inhibitors, 1-methyl-tryptophan (1-MT) and norharmane^25^. Corroborating the genetic data, both IDO1 inhibitors transiently reduced SNI-induced mechanical allodynia in a dose-dependent manner (Fig. 2h,i). Continuous high dose of 1-Mt treatment for one week (twice a day from day 14 to 20 after SNI) blocked mechanical allodynia over the treatment period (**Supplementary Fig. 2a**). Importantly, mechanical allodynia returned to the same level of SNI control group 2 days after the suspension of 1-MT treatment (Fig. 1e), indicating that IDO1 inhibition relieved but did not resolve mechanical allodynia (**Supplementary Fig. 2a**). Furthermore, a single treatment of mice with 1-MT at 21 days after SNI induction also reduced mechanical allodynia (**Supplementary Fig. 2b**). It is noteworthy that a potential role for the IDO1 in the activation of microglial cells in the spinal cord (ipsilateral dorsal horn) after peripheral nerve injury was ruled out since *Aif1* (IBA-1 gene) and *Cx3cr1* expression (marker of microglial cell activation) was comparable in WT and *Ido1*^-/-^ mice over a time course after SNI (**Supplementary Fig. 3a, b, d**). Noteworthy, we did not find any change in the marker of astrocytes activation (glial fibrillary acidic protein, *Gfap*) in the ipsilateral dorsal horn of the spinal cord after SNI even in the WT mice **(Supplementary Fig. 3c**). Collectively, these data indicate that IDO1 is involved in the maintenance of neuropathic pain but has no participation in nociceptive or inflammatory pain. Furthermore, they indicate that IDO1-derived kynurenines have no participation in microglial cell activation in the dorsal horn of the spinal cord after peripheral nerve injury.

**Fig. 2.**
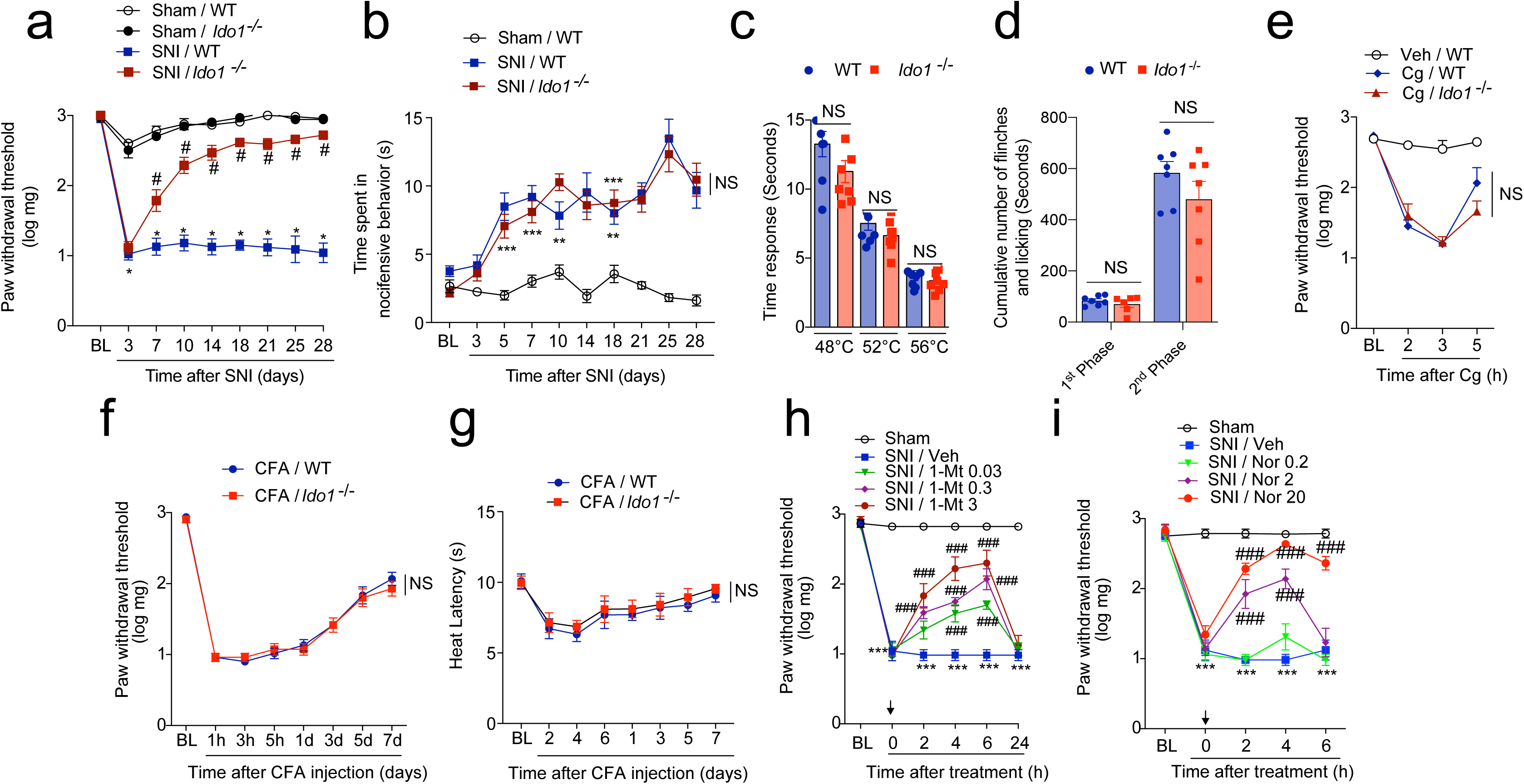
IDO1 is involved in the maintenance of neuropathic pain but has no role in nociceptive or inflammatory pain. **(a and b)** Mechanical and cold nociceptive responses were evaluated before and up to 28 days after SNI and sham surgeries in WT and *Ido1*^-/-^ mice. (n=9). **(c)** The nociceptive thermal threshold was tested in naïve WT and *Ido1*^-/-^ at 48 °C, 52 °C and 56 °C using the hot-plate test (n=8) **(d)** Formalin 1% was used to produce overt pain-like behavior. Total duration (sec) of nociceptive behaviors for 0–10 min (1st phase) and for 10–50 min (2nd phase) after formalin injection was evaluated in WT and *Ido1*^-/-^(n=8) **(e)** Mechanical nociceptive threshold using von Frey filament (n=4-5) was evaluated in WT and *Ido1*^-/-^ mice followed by intraplantar injection of carrageenan (Cg-100 µg per paw) or vehicle (saline). **(f and g)** Mechanical and thermal (heat) nociceptive threshold using von Frey filament (n=7) and Hargreaves’ test, respectively, were evaluated in WT and *Ido1*^-/-^ mice followed by intraplantar injection of CFA (10 µl per paw). **(h and i)** Mechanical nociceptive threshold was determined before and 14 days after SNI. Mice were treated i.p. with vehicle or 1-methil-DL-tryptophan (1-Mt, 0.03 - 3 mg/per mouse) or norharmane (Nor, 0.2 - 20 mg/kg) and mechanical allodynia was measured up to 24 h after treatment (n=6). Data are expressed as the mean ± s.e.m. \**P* < 0.05, \*\**P* < 0.01, \*\*\**P* < 0.001 versus sham or vehicle. # *P* < 0.05, ###*P* < 0.001 versus *Ido1^-/-^* mice and NS = no statistical significance. Two-way ANOVA, Bonferroni’s post-test (a and b, e-i); Unpaired Student’s *t-*test (c and d).

### IDO1 expressed in hematopoietic cells mediates neuropathic pain development

Despite indications of IDO1 protein expression in spinal cord from western blot analyses (Fig. 1e), no significant immunostaining for IDO1 was observed in the spinal cord after SNI (data not shown). However, perfusing (PBS) mice before spinal cord harvesting, eliminated IDO1 protein expression in the spinal cord as assessed by western blot (Fig. 3a). This suggested that IDO1, which is playing a role in the maintenance of neuropathic pain, is not expressed (upregulated) in the spinal cord resident cells, rather it is within the vasculature in the circulating (blood) immune cells. In line with this hypothesis, systemic plasma levels of Kyn increased after SNI and peaked 14 days after peripheral nerve injury (Fig. 3b). Corroborating the idea that IDO1 is up regulated in leukocytes after peripheral nerve injury, IDO1 expression (mRNA and protein) increased in the draining lymph nodes (dLNs) correspondent to nerve injury (popliteal and inguinal) (Fig. 3c and d) but not in the spleen after SNI surgery (Fig. 3e). To test if IDO1 expression in immune cells is important for the maintenance of neuropathic pain, bone marrow (BM) chimeric mice were generated. Irradiated *Ido1*^-/-^ mice that received BM from *Ido1*^-/-^ mice were still resistant to SNI-induced mechanical allodynia when compared to irradiated WT mice that received BM from WT mice (Fig. 3f). Adoptive transfer of BM from WT mice to irradiated *Ido1*^-/-^ mice restored mechanical allodynia of irradiated WT mice that received WT BM (Fig. 3f). On the other hand, irradiated WT mice receiving BM from *Ido1*^-/-^ mice became resistant to SNI-induced mechanical allodynia (Fig. 3f). These results indicate that IDO1 expressed by peripheral hematopoietic cells is essential to maintain neuropathic pain after peripheral nerve injury.

**Fig. 3.**
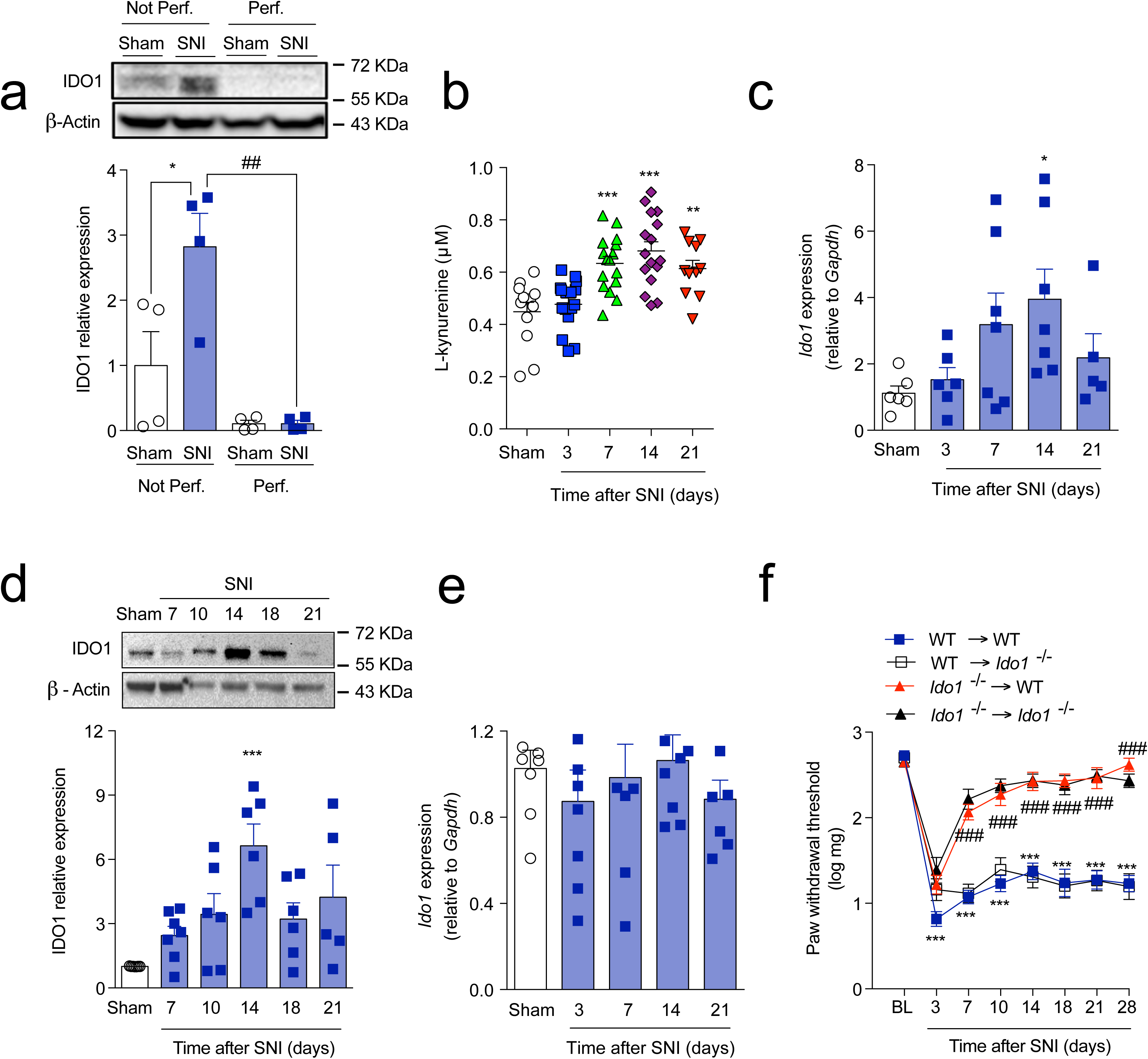
Up-regulation of IDO1 in peripheral hematopoietic cells compartment mediates neuropathic pain. **(a)** Western blotting analysis of IDO1 expression in the dorsal horn of the spinal cord with or without transcardiac perfusion with PBS after sham or SNI surgeries (n=4). **(b)** Time-course of kynurenine (Kyn) levels in the serum of mice after sham (14 days) and SNI surgeries (n=11-17 per time point). Time-course of *Ido1* mRNA **(c)** (n=5-8 per time point) and protein **(d)** (n=7 per time point) expressions in the draining lymph nodes after sham or SNI surgery. **(e)** Timecourse of *Ido1* mRNA expression in spleen after sham (14 days) or SNI surgeries (n=7-8 per time point). **(f)** Mechanical nociceptive threshold of WT→WT, *Ido1*^-/-^ →*Ido1*^-/-^, *Ido1*^-/-^ →WT and WT→ *Ido1*^-/-^ chimeric mice before (BL) and up to 28 d after SNI (n=8). Data are expressed as the mean ± s.e.m. \**P* < 0.05, \*\**P* < 0.01, \*\*\**P* < 0.001 versus sham or ##*P* < 0.01, ###*P* < 0.001 versus *Ido1^-/-^* mice or perfused group. Two-way ANOVA, Bonferroni’s post-test (f); One-way ANOVA, Bonferroni’s post-test (a-e).

### DCs-expressing IDO1 mediates neuropathic pain

Next, we sought to identify immune cells expressing IDO1 that contribute to the maintenance of neuropathic pain. We and others have identified dendritic cells (DCs) as the main immune cell population expressing IDO1 in many pathological conditions such as cancer, leukemia, arthritis, viral and bacterial infections^26–29^. Thus, we hypothesized that dendritic cells would be the cellular source of IDO1 that mediates neuropathic pain. To test this hypothesis, several approaches were employed. First, SNI was induced in CD11c^Yfp^ mice and CD11c^+^ cells (DCs) were isolated from dLNs by FACS sorting (**Supplementary Fig. 4**). Importantly, it was found that *Ido1* expression is higher in CD11c^+^ cells when compared to CD11c^-^ cells in sham operated mice (Fig. 4a). Moreover, SNI-induced upregulation of *Ido1* was only observed in CD11c^+^, but not in CD11c^-^ cells (Fig. 4a). Further addressing the importance of DCs for neuropathic pain maintenance, transgenic mice expressing the diphtheria toxin receptor (DTR) under the control of the CD11c (*Itgax*) promoter (CD11c-DTR-eGFP mice) were used to allow conditional depletion of DCs. As CD11c-DTR mice are sensitive to diphtheria toxin (Dtx) due to adverse effects causing fatal fulminant myocarditis^30^, we reconstituted lethally irradiated WT mice with BM cells from CD11c-DTR mice (donor) to generate mice that present DTR only in CD11c^+^ cells of the hematopoietic compartment (CD11c^DTR/hema^ mice, Fig. 4b). Following reconstitution, CD11c^+^ DCs can be depleted in these mice by the administration of Dtx ^25^. No adverse effects are associated with the chronic administration of Dtx to CD11c^DTR/hema^ chimeric mice^26^. Two months after BM reconstitution, SNI or sham surgery was performed in CD11c^DTR/hema^ mice followed by the treatment with Dtx or vehicle. Conditional depletion of DCs in CD11c^DTR/hema^ by Dtx treatment significantly reduced mechanical allodynia after SNI (Fig. 4c). FACS analysis of dLNs confirmed that Dtx treatments depleted CD11c^+^ DCs (Fig. 4d). Importantly, Dtx treatment had no effect on mechanical threshold or mechanical allodynia in WT mice either after sham or SNI surgery, respectively (**Supplementary Fig. 5**). The population of CD11c^+^ DCs in draining lymph nodes increased after SNI compared to sham operated mice (Fig. 4d), suggesting that nerve injury *per se* incited greater influx of DCs into dLNs. Additionally, SNI-induced IDO1 up-regulation was not observed in dLNs of DC-depleted mice (Fig. 4e). To further confirm that DCs-expressing IDO1 are important for the maintenance of neuropathic pain caused by peripheral nerve injury, *in vitro* differentiated BMDCs were generated and transferred (Fig. 4f)^31^. Whereas *Ido1*^-/-^ mice are resistant to SNI-induced mechanical allodynia, *Ido1*^-/-^ mice that received the transference of WT BMDCs became susceptible (Fig. 4g). Collectively, these results indicate that IDO1 is up regulated in periphery DCs and plays an important role in the maintenance of neuropathic pain after peripheral nerve injury.

**Fig. 4.**
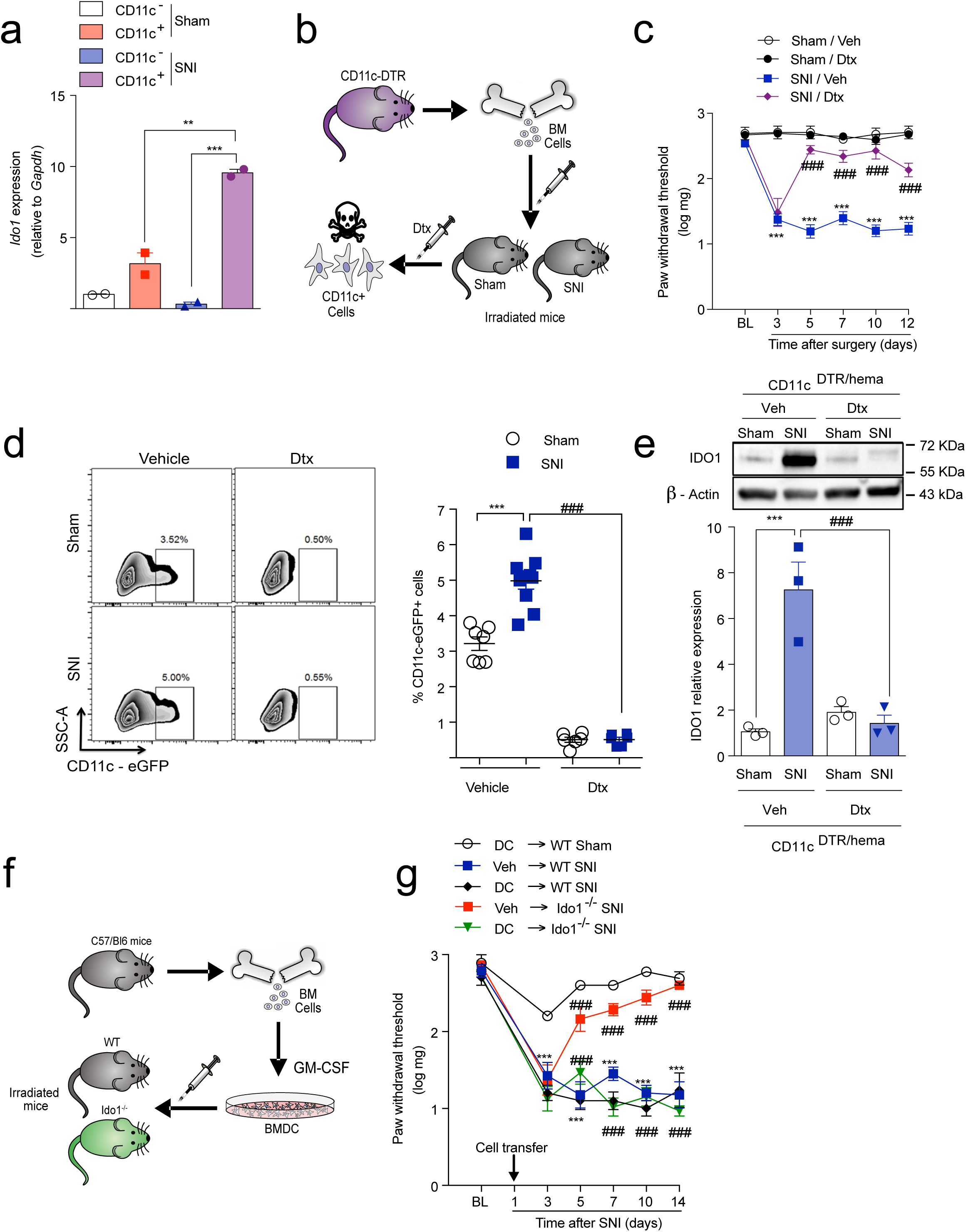
Dendritic cells-expressing IDO1 contributes to the maintenance of neuropathic pain. **(a)** *Ido1* mRNA expression in CD11c^+^ or CD11c^-^ cells isolated from the draining lymph nodes 14 days after sham or SNI surgeries (n=2 pooled 5). **(b)** Representative scheme of chimeric CD11c^DTR/hema^ mice establishment. **(c)** Mechanical nociceptive threshold was determined before and up to 12 days after sham or SNI surgeries in chimeric CD11c^DTR/hema^ mice treated with vehicle or diphtheria toxin (Dtx - 16 ng DTx/g; i.p.) (n=6). **(d)** Representative dot plots and quantification (percentage) of CD11c^+^ DC cells in the draining lymph nodes harvested 14 days after sham or SNI surgeries from chimeric CD11c^DTR/hema^ mice treated with vehicle or Dtx (n=6-10). **(e)** Western blotting analysis of IDO1 expression in the draining lymph nodes harvested 14 days after sham or SNI surgeries form chimeric CD11c^DTR/hema^ mice treated with vehicle or Dtx (n=3). **(f)** Representative scheme of dendritic cells differentiation and transfer to *Ido1*^-/-^ or WT mice. **(g)** Mechanical nociceptive threshold before (BL) and up to 14 days after sham or SNI in WT and *Ido1*^-/-^ that were transferred with *in vitro* differentiated DCs 1 day after surgeries (n=5-6). \**P* < 0.05, \*\**P* < 0.01, \*\*\**P* < 0.001 versus sham or ###*P* < 0.001 versus *Ido1^-/-^* mice or treatment. Data are expressed as the mean ± s.e.m. Two-way ANOVA, Bonferroni’s post-test (c and g); One-way ANOVA, Bonferroni’s post-test (a, d and e).

### DCs-expressing IDO1 accumulate into DRL and mediates neuropathic pain

There is evidence in the literature that peripheral immune cells (e.g. T cells) fail to infiltrate the spinal cord after peripheral nerve injury, but they might infiltrate/accumulate into the DRL of the somatosensory pathways^32–34^. These DRL-infiltrating leukocytes, specially CD4^+^ αβ T cells, play a role in the development of neuropathic pain^32^. After peripheral nerve injury, there is also an accumulation of CD45^+^ cells, specially macrophages into the dorsal root ganglion (DRGs)^35, 36^. Therefore, next, we sought to investigate whether DCs-expressing IDO1 would be also able to infiltrate/accumulate into the DRL or even the DRGs. Initially, we harvested DRGs (L3-L5) together with corresponding DRL and analyzed the expression of IDO1 after SNI induction. Western blotting analyses showed that the expression of IDO1 is upregulated in these tissues, peaking at 14 days after SNI induction (Fig. 5a). We then went on to assess the localization of IDO1 in these tissues. IDO1-expressing cells were found to increase in the vicinity of DRGs, which was assumed the DRL, but there were no obvious IDO1-expressing cells in the parenchyma of L4-5 DRGs (Fig. 5b,c). Since IDO1-expressing cells are found in the DRL after SNI and leukocytes in the DRL might influence spinal cord processes^32^, we sought to analyses whether its downstream metabolite (e.g kyn) would be also increased in the spinal cord. Mass spectrometry analyses of the dorsal horn of spinal cord tissue homogenate revealed an increase in the levels of Kyn in a time- and IDO-1-dependent manner (Fig. 5d**, Supplementary Fig. S6a**). Notably, Kyn levels in the spinal cord peaked 14 days after SNI, corroborating with the peak of IDO1 expression in the DRLs (Fig. 5d). In order to investigate whether IDO1 expressed in accumulated cells in the DRL might participate in the maintenance of neuropathic pain, SNI animals (14 days after injury) where intrathecally treated with IDO1 inhibitor. Notably, SNI-induced mechanical allodynia was inhibited by intrathecal treatment with 1-MT in a dose-dependent manner (Fig. 5e). In addition, the intrathecal treatment of SNI animals with 1-MT also reduced the levels of Kyn in the spinal cord (**Supplementary Fig. S6b)**, indicating that IDO-1-expressing cells in the DRL are probably the source of this KYNPATH metabolite detected in the spinal cord.

**Fig. 5.**
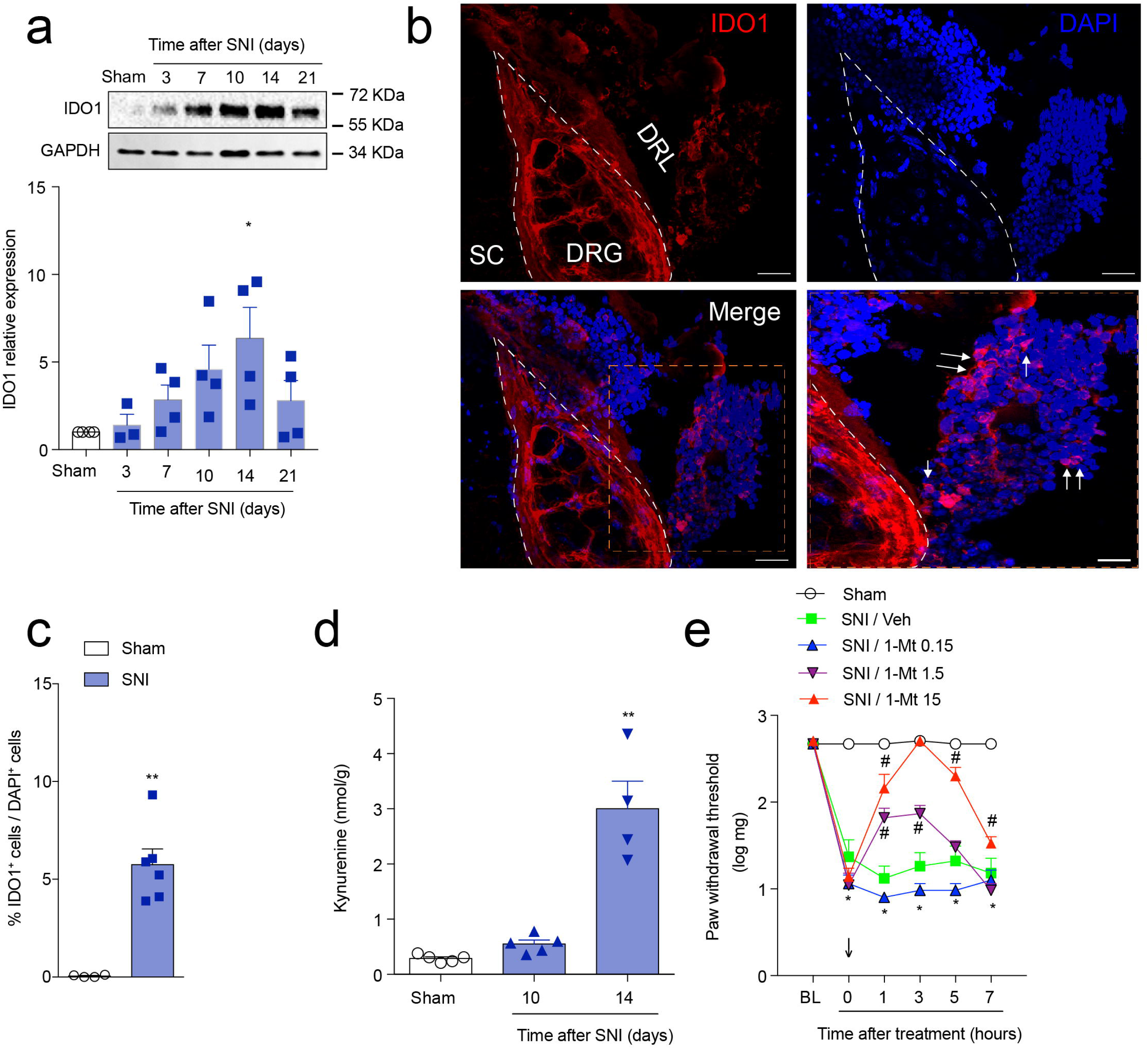
Cells expressing IDO1 accumulate in the DRL after SNI and contribute to the maintenance of neuropathic pain. **(a)** Time-course of IDO1 expressions in the DRGs plus DRL tissues (n=4 per time point). **(b)** Representative images and **(c)** quantification showing immunoreactivity for IDO1 (red color) double labeled with DAPI (cell nuclei; blue) in the ipsilateral region containing DRG (L4), DRL and spinal cord from SNI mice (14 day after SNI), bar scale 100 µm. **(c)** Quantification of IDO1-expressing cells in the DRL from SNI mice (14 day after SNI) or sham mice (n=6). **(d)** Time-course of kynurenine (Kyn) levels in the ipsilateral dorsal horn of the spinal cord of mice after sham (14 days) and SNI surgeries (n=4 per time point). **(e)** Mechanical nociceptive threshold was determined before and 14 days after SNI. Mice were treated intrathecally with vehicle or 1-methil-DL-tryptophan (1-Mt, 0.15 - 15 µg/per site) and mechanical allodynia was measured up to 7 h after treatment (n=5). Data are expressed as the mean ± s.e.m. *P < 0.05, **P < 0.001 versus sham group. # P < 0.05 versus vehicle treated mice. Two-way ANOVA, Bonferroni’s post-test (e); One-way ANOVA, Bonferroni’s post-test (a and d); Unpaired Student’s t-test (c).

Following we wanted to evaluate the cell subtype that express IDO-1 in the DRL after peripheral nerve injury. Based on our previous data, DCs were our immediately hypothesis. Initially, FACS analyses revealed that there is also an increase in CD11c+ cells in the DRL, 14 days after SNI induction (Fig. 6a). Remarkably, IDO1-expressing cells that accumulate in the DRL after SNI induction also express CD11c (Fig 6b). Notably, IDO-1-expressing cells in the DRL did not express a classical marker of macrophages (IBA-1, Fig. 6c). These results might confirm that CD11c^+^ cells-expressing IDO-1 that accumulate into DRLs after SNI are DCs. In attempt to eliminate and confirm the participation of CD11c-expressing IDO1 in the DRLs for the maintenance of neuropathic pain, SNI model was induced in CD11c-DTR mice. After 13 days, these mice were intrathecally treated with DTx or vehicle and mechanical allodynia was determined followed by the examination of IDO1 expression. We found that this treatment reduced SNI-induced mechanical allodynia (Fig. 6d), which was associated with a reduction of IDO1 expression in DRLs (Fig. 6e). Collectively, these results indicate that DCs-expressing IDO1 accumulate in the DRLs and mediates the maintenance of neuropathic pain.

**Fig. 6.**
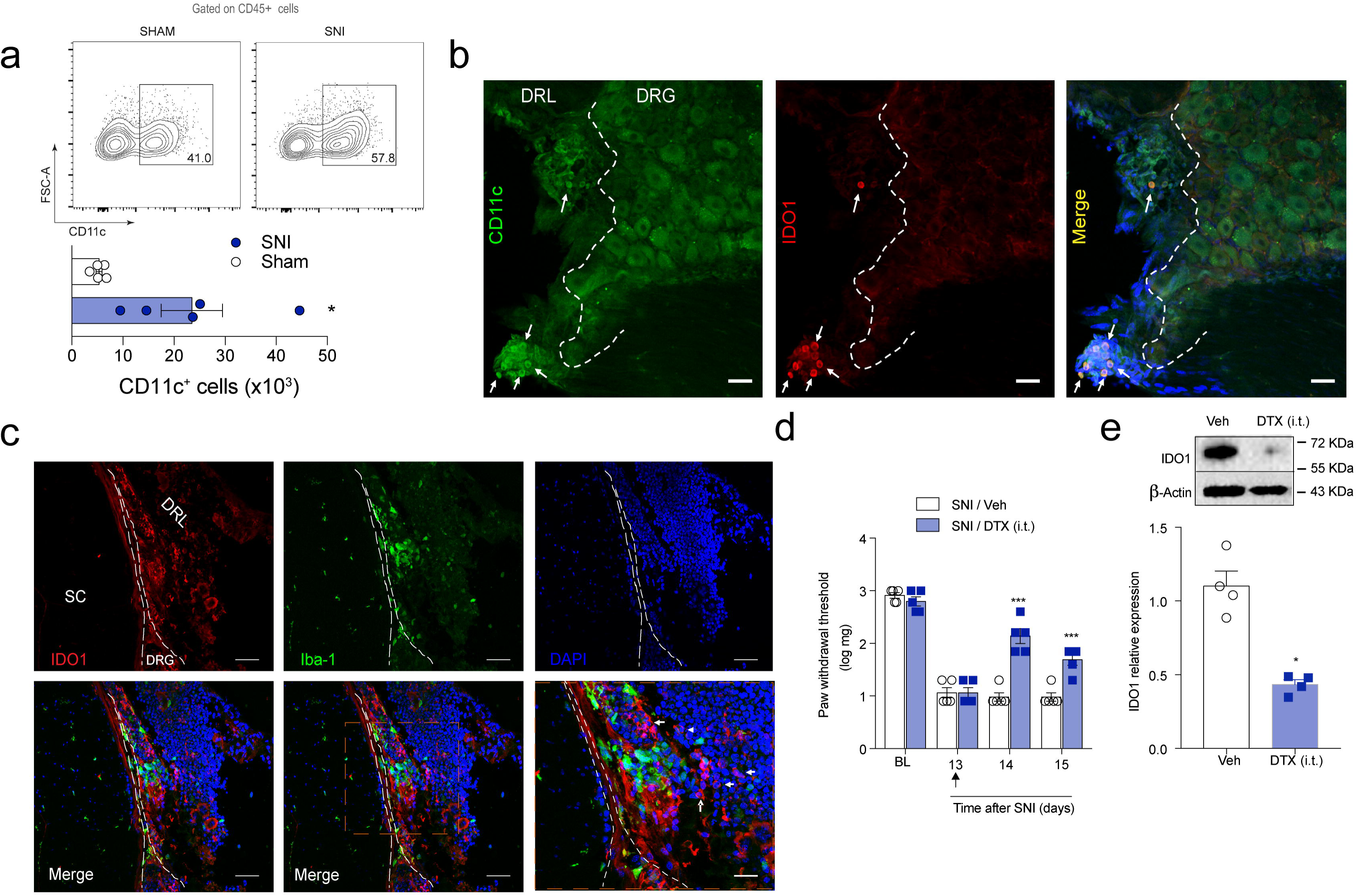
DCs-expressing IDO1 accumulate in the DRL after SNI and contributes to the maintenance of neuropathic pain. **(a)** Representative dot plots and quantification of CD11c^+^ (DCs) cells in DRGs plus DRLs harvested 14 days after sham or SNI surgeries from WT mice (n=5). **(b)** Representative images showing immunoreactivity for IDO1 (red color) double labeled with CD11c-eGFP (DCs) in the ipsilateral DRGs (L4) from SNI mice (14 day after SNI), bar scale 100 µm. **(c)** Representative images showing immunoreactivity for IDO1 (red color) double labeled with IBA-1 (macrophages) in the ipsilateral region containing DRG (L4), DRL and spinal cord from SNI mice (14 day after SNI), bar scale 100 µm. **(d)** Mechanical nociceptive threshold was determined before and 13 days after SNI. Mice were treated intrathecally with vehicle or Diphtheria toxin (DTx, 20 µg/per site) and mechanical allodynia was measured up to 48 h after treatment (n=5). **(e)** Western blotting analysis of IDO1 expression in the DRGs plus DRL tissues harvested 15 days after SNI mice treated with vehicle or Dtx (n=4). Data are expressed as the mean ± s.e.m. *P < 0.05, ***P < 0.001 versus sham group. # P < 0.05 versus vehicle treated mice. One-way ANOVA, Bonferroni’s post-test (d); Unpaired Student’s t-test (a, e).

### Spinal astrocytes expressing KMO mediates the maintenance of neuropathic pain

Next, we investigated the mechanisms by which IDO1 activity in the DRL maintains neuropathic pain. As we showed before, the accumulation of DCs-expressing IDO-1 in the DRLs rendered increased levels of Kyn in the spinal cord after peripheral nerve injury. So, next we investigated whether this spinal cord Kyn would be directly involved in the maintenance of neuropathic pain. In addition, as Kyn in the CNS is rapidly converted into downstream metabolites^37^, we might also hypothesized that Kyn that reached spinal cord after peripheral nerve injury could be converted into downstream neuroactive metabolites (e.g. 3-Hk and 3-Haa) and these metabolites directly account for the maintenance of neuropathic pain. To gain further information about these possibilities, initially we tested the ability of Kyn and its immediate downstream metabolites, 3-Hk and 3-Haa to produce mechanical pain hypersensitivity when injected intrathecally in naïve animals. When equimolar doses of these molecules where injected, it was found that only 3-Hk produced robust mechanical pain hypersensitivity (Fig. 7a **and Supplementary Fig. 7**).

**Fig. 7.**
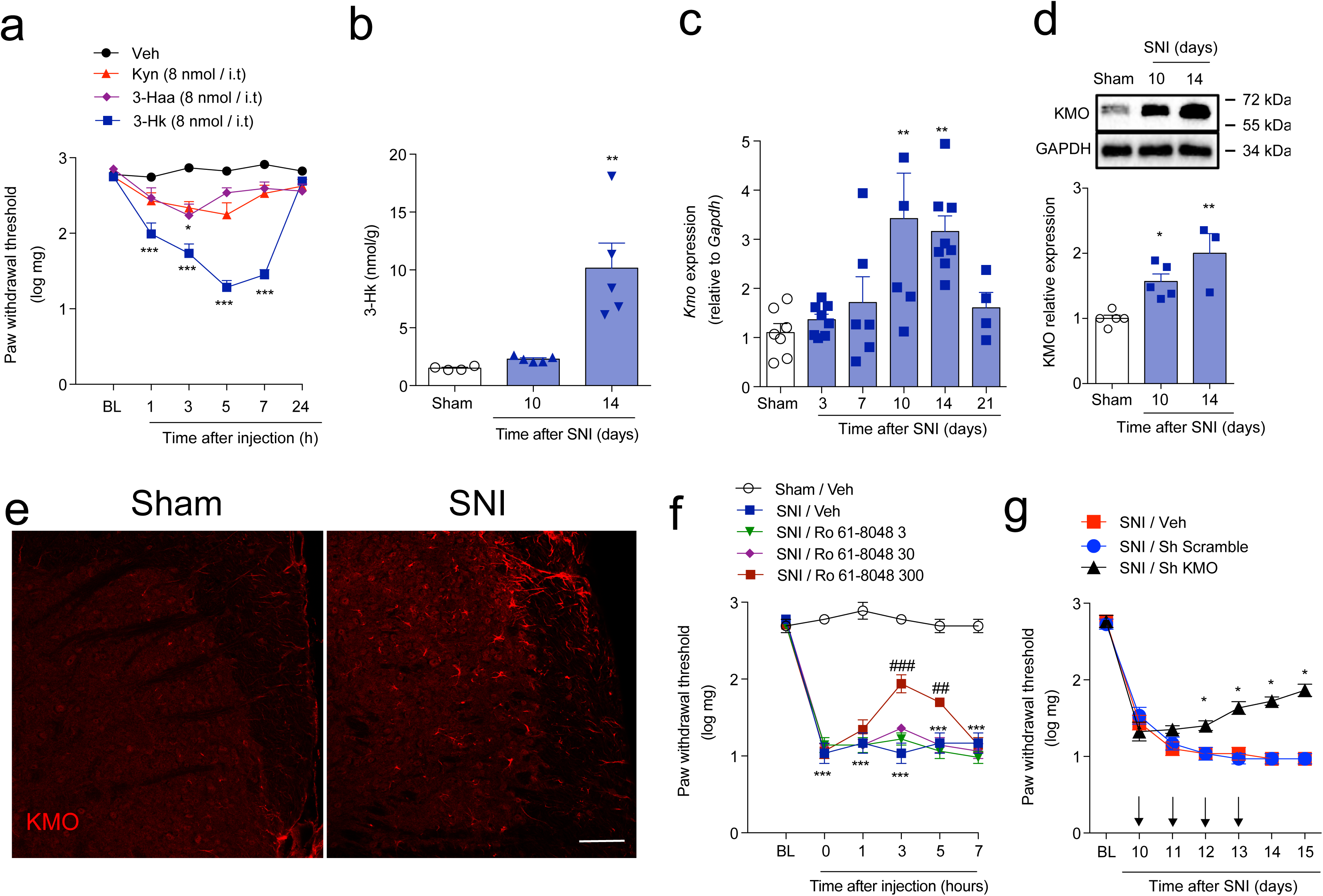
KMO is upregulated in the spinal cord after SNI and mediates neuropathic pain. (a) Mechanical nociceptive threshold was evaluated before and up to 24 h after intrathecal injection of equimolar dose (8 nmol) of kynurenine (Kyn), 3-hydroxykynurenine (3-Hk), 3-Hydroxyanthranilic acid (3-Haa) or vehicle (saline) in WT mice (n=4-6). This panel shows representative curves obtained in the dose-response analysis of mechanical allodynia caused by equimolar doses of Kyn, 3-Hk, 3-Haa in WT mice (Supplementary Fig. 8). (b) Time-course of 3-hydroxykynurenine (3-Hk) levels in the ipsilateral dorsal horn of the spinal cord of mice after sham (14 days) and SNI surgeries (n=4-5 per time point). (b) Time-course of *Kmo* mRNA expression in the ipsilateral dorsal horn of the spinal cord spinal cord after sham (14 days) and SNI surgeries. (n=5-8). (c) Western blotting analysis of KMO expression in the ipsilateral dorsal horn of the spinal cord after sham (14 days) or SNI surgeries (10 and 14 days). (n=4-5). Representative image of KMO expression analyzed by immunofluorescence in the dorsal horn of the spinal cord after sham or SNI induction (14 days). Scale bar 50 µm. (d) Mechanical nociceptive threshold was determined before and 14 days after SNI. Mice were then treated intrathecally (i.t.) with Ro 61-8048 (KMO inhibitor; 3–300 nmol) or vehicle and mechanical allodynia was measured up to 7 h after treatments (n=5). (d) Mechanical nociceptive threshold was determined before and 10 days after SNI followed by intrathecal treatment with ShRNA against KMO, ShRNA scramble or vehicle (indicated arrows) and mechanical allodynia was measured up to 15 days after SNI (n=6). Data are expressed as the mean ± s.e.m. *P < 0.05, **P < 0.01, ***P < 0.001 versus sham or saline injected. ##P < 0.01, ###P < 0.001 versus mice treated with Ro61-8048 or scramble ShRNA. Two-way ANOVA, Bonferroni’s post-test (a, f and g); One-way ANOVA, Bonferroni’s post-test (b, c and d).

The results showing that 3-Hk possess more potent pronociceptive activity than Kyn lead us to consider that IDO1 downstream metabolites other that Kyn (e.g. 3-Hk) would be more important for the maintenance of neuropathic pain. Because KMO is the rate-limiting downstream enzyme in the KYNPATH that oxidatively metabolizes Kyn into 3-Hk^4^, next we hypothesized that Kyn that reaches spinal cord might be metabolized by KMO to generate downstream pronociceptive factors, such as 3-Hk, which might be involved in the maintenance of neuropathic pain. Consistent with this hypothesis, KMO downstream metabolite, 3-Hk, levels increased in the ipsilateral dorsal horn of the spinal cord after SNI induction (Fig. 7b), which was dependent on IDO1 activity in the DRLs (**Supplementary Fig. S6c)**. This was also associated with an increase in KMO expression (mRNA and protein) in the ipsilateral dorsal horn of the spinal cord after SNI induction (Fig. 7c-e). Notably, no significant changes in the levels of Kyna were observed in the spinal cord after SNI, although the ratio 3-Hk/Kyna increased significantly (**Supplementary Fig. 8 a and b).** Additionally, we found that pharmacological treatment (intrathecally) with KMO inhibitor Ro 61-8048^38^ also reduced SNI-induced mechanical allodynia in a dose-dependent manner (Fig. 7f). Furthermore, intrathecal treatment with shRNA against KMO also reduced SNI-induced mechanical allodynia (Fig. 7g).

In order to characterize which spinal cord cell subtype might be expressing KMO and would be important for the production of 3-Hk, a series of experiments were performed. Firstly, immunofluorescence of spinal cord slices from SNI animals, revelated that KMO expression was detected exclusively in glial fibrillary acidic protein (GFAP)^+^ cells (Fig. 8a**; Supplementary Fig. 9, Movie 1**) but not in IBA-1^+^ microglia (Fig. 8a**; Supplementary Fig. 9, Movie 2**) nor NeuN^+^ neurons (Fig. 8a**; Supplementary Fig. 9, Movie 3**), indicating selective KMO expression by astrocytes, which are in close contact with spinal cord neurons (**Movie 3**). Next we performed cells sorting analyses of microglia and astrocytes from spinal cord^39, 40^ followed by the analyses of *Cx3cr1*, *Gfap* and *Kmo* expression (Fig. 8c-e). Supporting the immunofluorescence data, we found that *Kmo* mRNA expression is higher in astrocytes than microglial cells (Fig. 8e). To further support these data, pure primary astrocytes from newborn mouse cortex (**Supplementary Fig. 10**) were culture and the expression of KMO was analyzed. Activation of primary culture astrocytes with TNF or microglia conditional medium (MCM) induced an upregulation in the expression (mRNA and protein) of KMO (Fig. 9a-c). Importantly, activation of differentiated U87-MG, an immortalized human astrocytic cell line^41^, induced the increase in the expression of KMO (Fig. 9d). Finally, we sought to confirm whether KMO expressed in spinal cord astrocytes is important for the maintenance of neuropathic pain. For that, we knocked down the expression of KMO specifically in astrocytes using a lentivirus-delivered shRNA (shKMO) expressed under the control of the *Gfap* promoter (Fig. 9e)^42, 43^. The lentivirus was administered intraspinal^44^ 10 d and 13 days after SNI induction; a lentivirus carrying a nontargeting shRNA was used as a control (sh scramble). KMO silencing in spinal cord astrocytes (Fig. 9f) reduced SNI-induced mechanical allodynia (Fig. 9g). Collectively, these results indicated that after peripheral nerve injury astrocytes expressing KMO mediates the maintenance of neuropathic pain probably through conversion of Kyn into downstream pronociceptive factors, such as 3-Hk.

**Fig. 8.**
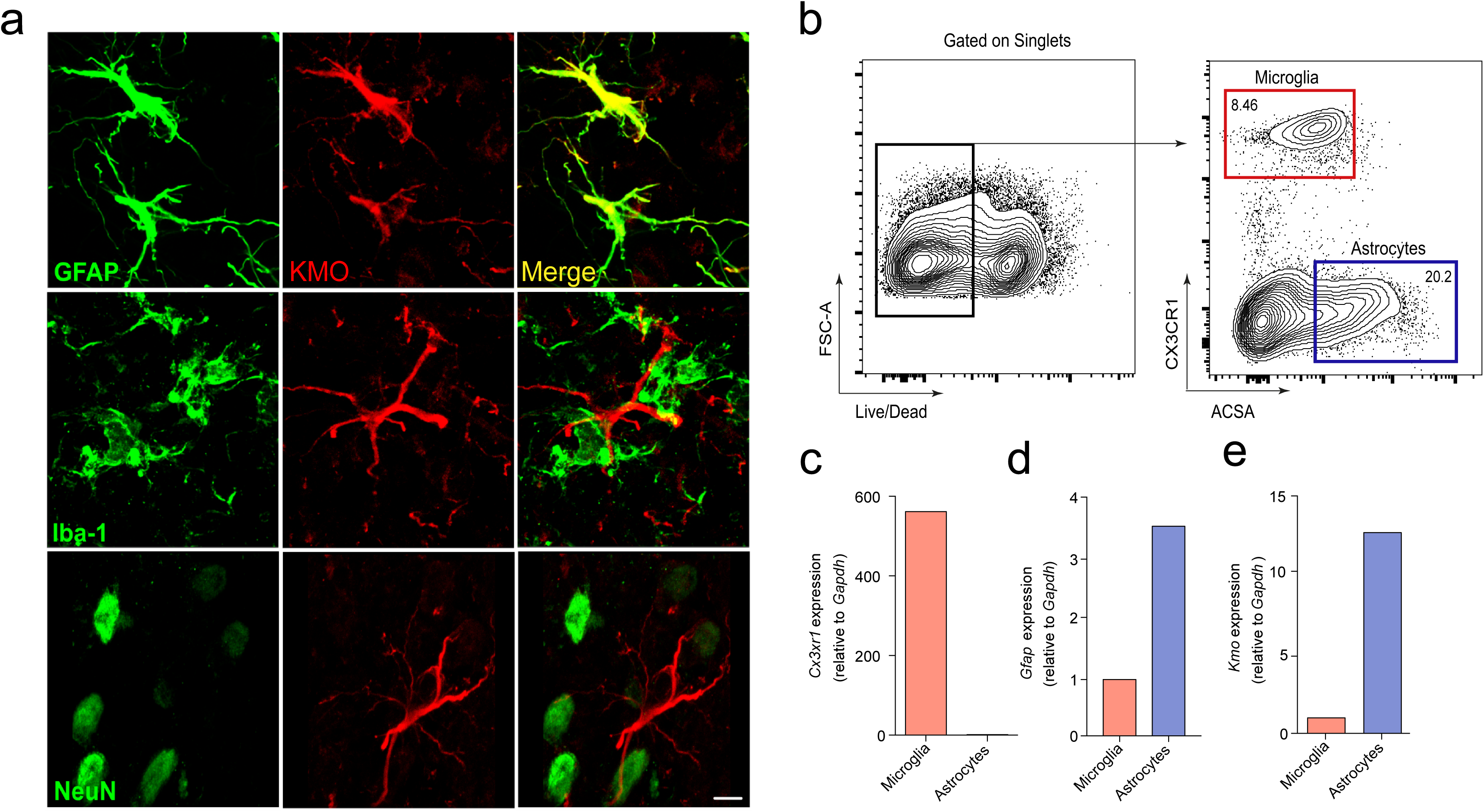
KMO expression cell profile in the spinal cord after SNI. (a) Representative images showing immunoreactivity for KMO (red color) double labeled with anti-GFAP (astrocytes), anti-IBA1 (microglia) or anti-NeuN (neurons) in the ipsilateral dorsal horn of the spinal cord from SNI mice (14 day after SNI), bar scale 5 µm. (b) Representative FACS sorting strategy for microglia cells (CX3CR1-eGFP+ cells) and astrocytes (ACSA2+) isolation from spinal cord (b) Cx3cr1, Gfap and Kmo mRNA expression in microglia and astrocytes isolated from the spinal cord (pooled of 8).

**Fig. 9.**
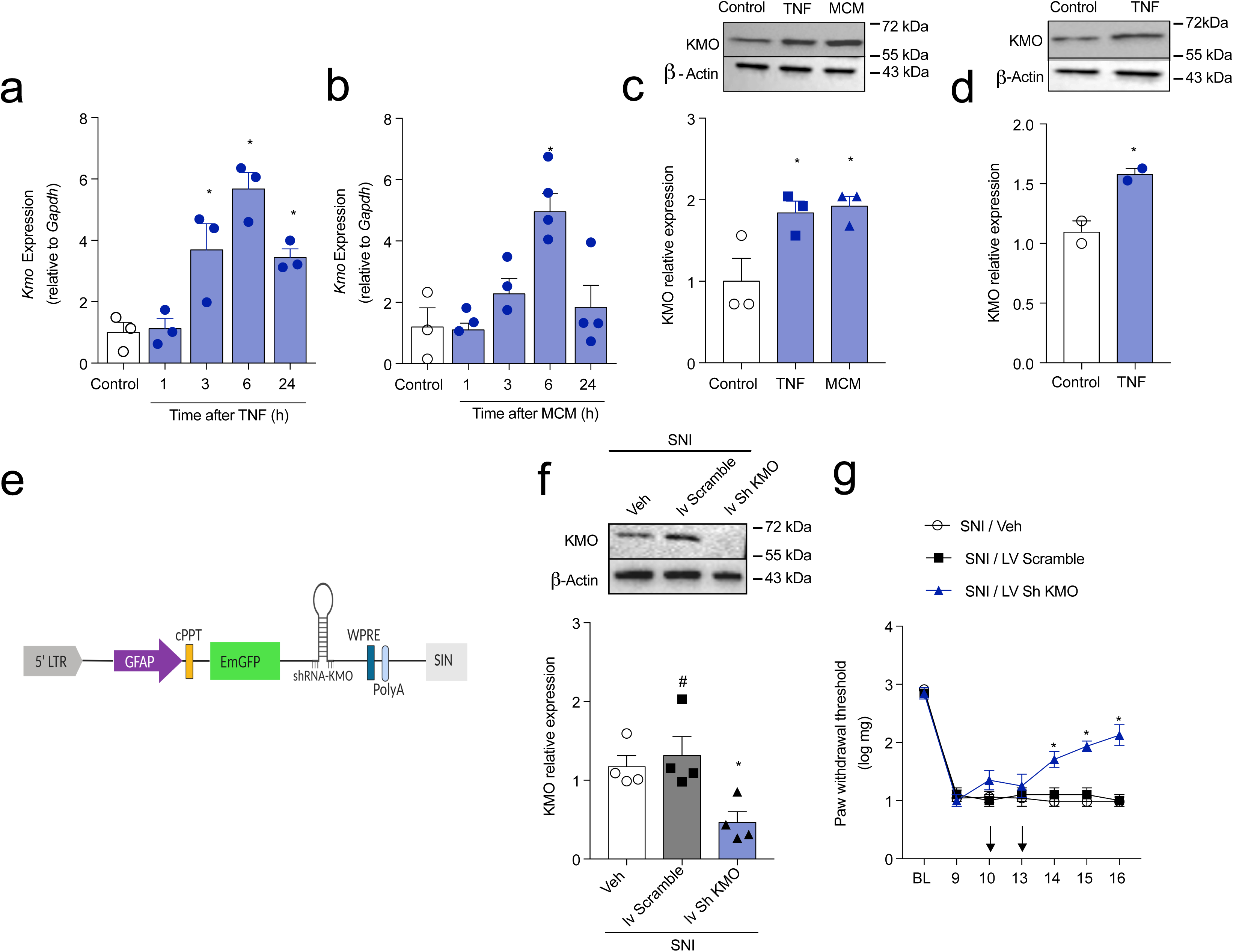
Astrocytes-expressed KMO maintains neuropathic pain. Primary cultured astrocytes from mouse cortex were stimulated with **(a)** TNF (10 ng/ml) and **(b)** microglia conditional medium (MCM). After indicated time points, mRNA was extracted and *Kmo* expression was analyzed by real-time PCR. **(c)** Primary cultured astrocytes from mouse cortex were stimulated with TNF (10 ng/ml) or MCM. The expression of KMO in the cell extract was evaluated 24 h after stimulation by western blotting. **(e)** Schematic of the astrocyte-specific shRNA targeting lentiviral vector used to knockdown KMO in astrocytes of the spinal cord. **(f)** Mechanical nociceptive threshold was determined before and 9 days after SNI followed by intrathecal treatment with lentiviral vectors expressing ShRNA Control or shRNA Kmo (n = 4-5) at days 10 and 13 after SNI. Mechanical allodynia was measured up to 16 days after SNI and ipsilateral dorsal horn of the spinal cord was collected for analyses of **(g)** KMO expression (n=4). Data are expressed as the mean ± s.e.m. \**P* < 0.05, versus medium treated # *P* < 0.05, versus mice treated with scramble ShRNA. Two-way ANOVA, Bonferroni’s post-test (f); One-way ANOVA, Bonferroni’s post-test (a-d and g).

## Discussion

Peripheral nerve injury-induced neuropathic pain development depends on complex multifactorial pathophysiological processes^1, 43, 45^. It is thought that the knowledge of the cellular and molecular mechanisms involved the transition from acute to chronic neuropathic pain after peripheral nerve injury would be the best way to discovery novel targets for its control and/or prevention^46, 47^. In this context, in the present study we investigated the potential role of KYNPATH in the development of neuropathic pain followed peripheral nerve injury. We found that pharmacological or genetic inhibition of crucial canonical enzymes of KYNPATH (IDO1 and KMO) reduced stablished neuropathic pain. We also found that after peripheral nerve injury there is an accumulation of DCs-expressing IDO1 into the DRLs which promotes an increase in the spinal cord levels of Kyn. Kyn is subsequently metabolized into a potent pro-nociceptive molecule, 3-Hk, by astrocytes-expressing KMO. These data indicate a novel important mechanism of neuronimmune-glia crosstalk in the pathophysiology of neuropathic pain induced by peripheral nerve injury.

The participation of KYNPATH in the development of pathological pain has been investigated^18–20, 27, 48^. For instance, IDO1 genetic deficiency and pharmacological inhibition alleviate arthritic pain^48^. Furthermore, we have also shown the participation of IDO1 in the genesis of virus infections-incite pain hypersensitivity^27^. Regarding neuropathic pain, the participation of KYNPATH, at least of IDO1, is controversial^18–20^. Whereas intrathecal treatment with 1-Mt reduced mechanical allodynia triggered by peripheral nerve injury in rats, supporting our findings, *Ido1*^-/-^ mice developed mechanical allodynia similar to WT mice^18, 20^, which is in contrast to our data. Although the explanation for this discrepancy is not immediately apparent, we can speculate some possibilities, such as animal colony with different microbiota, different method to quantify mechanical allodynia (up and down method *versus* absolute threshold). Nevertheless, similar to our findings none of these studies detected expression of IDO1 in the spinal cord after peripheral nerve injury^18, 20^, but they found IDO1 up-regulation in the DRGs in rats^20^ and systemic plasma increase of kyn levels in SNI mice^18^. SNI-induced neuropathic pain is mainly characterized by the development of mechanical and cold allodynia^49^. Interestingly, KYNPATH seems to be involved only in SNI-induced mechanical allodynia but not in cold allodynia. It is in line with recent evidence suggesting that neuro-immune interactions are important for SNI-induced mechanical allodynia whereas has no role in the development of cold allodynia^49^.

Here we provided additional evidence on the mechanisms by which IDO1 is up regulated in the DRGs after peripheral nerve injury. Indeed, we found that IDO1 is expressed in DCs that accumulate into DRL. Corroborating, accumulation of peripheral leukocytes into the DRL after peripheral nerve injury has been demonstrated^32, 50, 51^. For instance, CD4+ Th1 cells generated in the draining lymph nodes of the peripheral nerve injury accumulate in the DRL and drive glial and neuronal chances in the spinal cord that in turn mediate neuropathic pain development^32^. Thus, our study provides evidence that not only T-cells are able to accumulate in the DRLs, but also DCs. Although our data did not provide evidence about the exact origin of these IDO-1-expresssing DCs that accumulate into DRLs, the fact that the population of these cells also increased in the draining lymph nodes and the spinal cord circulatory system would indicate they might be generated peripherally and infiltrate into DRL. Nevertheless, we could not discard that cerebrospinal meninges resident DCs^52^ could also migrate toward DRL after nerve injury. Further studies are necessary to elucidate this point.

Another intriguing finding of our study was the role of KMO expressed by astrocytes in the maintenance of neuropathic pain. KMO expression has been identified mainly in neurons and microglial cells, at least in the brain^53–56^. Furthermore, the stimulation of primary culture of microglia with LPS, but not of astrocytes, increased the expression of KMO^19^. On the other hand, there is evidence that KMO is expressed in astrocytes of the spinal cord and of the forebrain^56^. Herein we used a considerable number of different approaches to show that in the spinal cord, KMO is mainly expressed and upregulated in astrocytes after peripheral nerve injury. Furthermore, the specific knockdown of KMO in spinal cord astrocytes was able to reduce mechanical allodynia triggered by peripheral nerve injury, indicating its role in the maintenance of neuropathic pain. Supporting the importance of KMO in maintenance of neuropathic pain, we also provided evidence that its main downstream metabolite, 3-Hk, is a very potent pro-nociceptive molecule. The fact that 3-Haa weakly produced mechanical allodynia when compared with its upstream precursor, 3-Hk, might indicate that 3-Hk is one of the final directly acting pronociceptive mediator. It is important to point out that targeting KMO, for instance with pharmacological inhibitor, besides to inhibit the formation of the pro-nociceptive 3-Hk, it could also divert KYNPATH into the formation of an antinociceptive molecule, KYNA^57^. In fact, KYNA has been considered a neuroprotective endogenous antagonist of NMDA with analgesic properties^57, 58^. Additionally, more recently KYNA has been also identified as an endogenous agonist of GPR35 which is also a possible alternative mechanism to control pathological pain^59–61^. Another important point that emerge from our results is regarding the possible mechanisms by which 3-Hk promotes pain hypersensitivity. Although the direct biding sites for 3-Hk are not known, several studies indicate its ability to trigger the formation of toxic free radicals, especially superoxide and hydrogen peroxide, mitochondrial damage that in turn leading to neuronal dysfunction^62–65^. Importantly, all these mechanisms are important for the development of neuropathic pain^66–72^. Therefore, we might suggest that the increase of KMO-dependent 3-Hk in the spinal cord after peripheral nerve injury would be a possible driver of oxidative stress in neuropathic pain development.

Neuropathic pain can be triggered by several different types of nerve injury or diseases of the somatosensory nervous system^1, 73, 74^. It varies from chronic pain caused by physical trauma, as has been mimicked here (SNI model), up to metabolic disorders (eg. Diabetes), infection (eg. HIV) and chemically-induced (e.g. cancer chemotherapy)^75^. Although there is some common pathophysiological mechanism involved in the development of these neuropathic pain subtypes, they also have specific not-shared mechanisms^45, 76–78^. Although we did not investigate the participation of KYNPATH in other models of neuropathic pain, there is evidence indicating an upregulation of KYNPATH in some diseases that lead to neuropathic pain development, such as diabetes^79^ and HIV^80^. For instance, there is an increased concentration of the 3-Hk in the CNS of HIV-1 patients^81^. In addition, KYNPATH enhancement is generally observed in different types of cancer, important clinical conditions, in which pain has a neuropathic component^82–84^. Therefore, further studies will be necessary to addresses whether KYNPATH is a common mechanism among different types of neuropathic pain.

In summary, in the present study we provided an ample characterization of the role of the KYNPATH in the maintenance of neuropathic pain. As depicted in the **Supplementary Fig. 11**, after a peripheral nerve injury, IDO1 is up regulated in peripheral DCs, which might be able to accumulate into DRL leading to an increase in the spinal cord levels of Kyn. Kyn is then metabolized locally by astrocytes-expressing KMO into a potent pronociceptive molecule 3-Hk. These data reveal a previously unappreciated role for the KYNPATH as a critical link between immune system (DCs), spinal cord glia cells (astrocytes) and neuronal sensitization to maintain neuropathic pain. These mechanisms offer new targets for drug development against this type of chronic pain.

## Methods

### Animals

The experiments were performed in C57BL/6 male mice (Wild Type, WT, 20-25 g) and mice deficient in the following protein: indoleamine 2,3 dyoxigenase type-1 (*Ido1*^-/-^)^85^. We generated chimeric mice using bone marrow (BM) cells from CD11c-DTR-eGFP transgenic mice, which express diphtheria toxin receptor and eGFP under control of *Itgax* gene promoter (Itgax encoding CD11c)^86^. We also used CD11c-eYFP mice that express enhanced yellow fluorescent protein (eYFP) under control of *Itgax* gene promoter^87^. Transgenic mice expressing the green fluorescent protein (GFP) in cells that express CX3C chemokine receptor 1 (CX3CR1^GFP/+^)^88^ were used for FACS sorting analyses of the microglia and astrocytes in the spinal cord. Local colonies of transgenic mice were then established and maintained on a C57BL/6 background at the animal care facility of Ribeirão Preto Medical School, University of São Paulo. Animals were taken to the testing room at least 1 h before experiments and were used only once. Food and water were available *ad libitum*. Animal care and handling procedures were in accordance with the International Association for the Study of Pain guidelines^S1^ for those animals used in pain research and they were approved by the Committee for Ethics in Animal Research of the Ribeirao Preto Medical School - USP (Process n° 045/2013 and 120/2018). Four to twelve randomly assigned mice were used per experimental group per experiment. Mice were selected based on their genotype. All other criteria were not considered and as such, randomized. We were not blinded to mice genotypes or group allocations except where it is mentioned.

### Neuropathic pain model (Spared nerve injury)

A model of persistent peripheral neuropathic pain was induced as previously described^24^. Under isoflurane (2%) anesthesia the skin on the lateral surface of the thigh was incised and a section made directly through the biceps femoris muscle exposing the sciatic nerve and its three terminal branches: the sural, common peroneal and tibial nerves. The Spared Nerve Injury (SNI) comprised an axotomy and ligation of the tibial and common peroneal nerves leaving the remaining sural nerve intact. The common peroneal and the tibial nerves were tightly ligated with 5.0 silk and sectioned distal to the ligation, removing 2 ± 4 mm of the distal nerve stump. Muscle and skin were closed in two layers. For sham-operated mice, the sciatic nerve and its branches were exposed without lesioning. Then, mechanical pain hypersensitivity was evaluated up to 21 days after the surgery.

### Inflammatory pain model

Mice received an injection of carrageenan (Cg-100 μg/paw), CFA (10μL) or vehicle (saline) subcutaneously into the plantar region of the right hindpaw^89, 90^. Then, mechanical^91^ and thermal (heat)^92^ pain hypersensitivity was determined at indicated time points after stimulus injection.

### Behavioral nociceptive tests

An investigator blinded to group allocation performed all the behaviors tests.

### Formalin nociception test

We assessed formalin-evoked nociception by injection of 20 µl of formalin (1%) into the dorsal surface of the right hind paw of mice. The time in seconds spent licking or flinching the injected paw was recorded and expressed as the total nociceptive behaviors in the early phase (0–10 min) and late phase (10–50 min) after the formalin injection^93^.

### Thermal Nociceptive Test

The latency of paw withdrawal to radiant heat stimuli was measured using a Plantar Ugo Basile apparatus (Stoelting), as previously described^92^. Mice can move freely in this apparatus on an elevated glass surface with plastic boxes above as the top cover. Mice were given a 1-h acclimation period before testing until they became calm and motionless. A calibrated infrared light source of high intensity was applied perpendicular on the plantar surface of each mouse’s hind paw. The end point was characterized by the removal of the paw followed by clear flinching movements. Latency to paw withdrawal was automatically recorded. Each hind paw was tested alternately with an interval of 5 min for four trials.

### von Frey filament test

For testing mechanical nociceptive threshold, mice were placed on an elevated wire grid and the plantar surface of the ipsilateral hind paw stimulated perpendicularly with a series of von Frey filaments (Stoelting, Chicago, IL, USA) with logarithmically increasing stiffness (0.008–2.0g). Each one of these filaments was applied for approximately 3-4s to induce a paw-withdrawal reflex. The weakest filament able to elicit a response was taken to be the mechanical withdrawal threshold. The log stiffness of the hairs is determined by log10 (milligrams) and ranged from 0.903 (8 mg or 0.008 g) to 3.0 (1000 mg or 1 g)^91, 94^.

### Hot-Plate Test

The noxious heat thresholds of the hind paws were also examined using the Hot-Plate test. Mice were placed in a 10-cm-wide glass cylinder on a hot plate (IITC Life Science) maintained at 48°C, 52°C or 56°C. Two control latencies at least 10 min apart were determined for each mouse. The latencies in seconds for paws licking or jumping for each animal were recorded. To minimize tissue damage, a maximum latency (cut-off) was set at 20 s^95^.

### Acetone test

Mice were placed in a clear plastic box with a wire mesh floor and allowed to habituate for 30 minutes prior to testing. Then, 50 μl fluid (acetone) was sprayed on the plantar surface of the hind right paw using a syringe of 1 ml (Tuberculin slip tip, BD, Franklin Lake, NJ, USA). Paw nociceptive responses, defined as flinching, licking or biting of the limb were measured within 1 minute after the application of acetone^96^.

### Drugs Administration

#### The i.p. administration

the drugs were injected into the ventral portion of the mouse close to the abdominal midline using a 1 mL syringe with a needle length of 2.5 cm and 0.7 cm gauge.

#### Intrathecal injection

The technique used for intrathecal injection was done as described previously^97^ with modifications. Under isoflurane (2%) anesthesia, mice were securely hold in one hand by the pelvic girdle and inserting a BD Ultra-Fine® (29G) insulin syringe (BD, Franklin Lakes, NJ, USA) directly on subarachnoid space (close to L4–L5 segments) of the spinal cord. A sudden lateral movement of the tail indicated proper placement of the needle in the intrathecal space. For all administrations was used 5 µl of volume. Then, the syringe was held in the specific position for a few seconds and progressively removed to avoid any outflow of the substances

### Enzymatic assay for IDO activity

IDO activity was assayed as described in a previous report^98^. In brief, spinal cord tissues (L4-L6 level) were homogenized with a Polytron homogenizer (Kinematica, Lucerne, Switzerland) in 1.5 volumes of ice-cold 0.14 M KCl-20 mM potassium phosphate buffer (pH 7). The homogenate samples were centrifuged at 7000 X g and 4°C for 10 min. An aliquot of supernatant was taken for the measurement of IDO activity. The reaction mixture contained 50 µl enzyme preparation and 50 µl substrate solution. The composition of the substrate solution was 100 mM potassium phosphate buffer (pH 6.5), 50 µM methylene blue, 20 µg catalase, 50 mM ascorbate, and 0.4 mM L-TRP. Post-incubation of the reaction mixture at 37°C, samples were acidified with 3% perchloric acid and centrifuged at 7000 X g and 4°C for 10 min. The concentrations of the enzymatic products were measured by using HPLC. Enzyme activity was expressed as the product content per hour per gram of tissue protein.

### Measurement of plasmatic l-kynurenine concentration

Plasma concentration of kynurenine (Kyn) was measured by high-performance liquid chromatography (HPLC) with a spectrophotometric detector (TOSOH UV-8000) or fluorescence spectrometric detector (HITACHI, Tokyo, Japan) as described previously^10^. Briefly, separation was obtained with a reverse-phase column (Brave ODS 3 μm 150 mm × 4.6 mm; Alltech, IL, U.S.A.) and a mobile phase (flow rate 0.75 mL/min) composed of 0.1 M sodium acetate, 0.1 M acetic acid and 1% acetonitrile. The fluorescence excitation and emission wavelengths were set at 270 and 360 nm, respectively. UV signals were monitored at 355 nm for Kyn.

### Measurement of tissue kynurenines concentration

#### Chemicals & reagents

L-Kynurenine (Kyn), 3-hydroxykynurenine (3-Hk), kynurenic acid (Kyna), 3-hydroxyanthranilic acid (3-Haa), ascorbic acid (AA), formic acid (FA) and trifluoroacetic acid (TFA) were obtained from Sigma Aldrich (Steinheim, Germany). Methanol (MeOH; LC–MS grade), acetonitrile (MeCN, LC–MS grade) were obtained from Merck KGaA (Darmstadt, Germany). The internal standard (IS) was: 6-Chloro-DL-tryptophan (TRP_Cl; Goldbio, St Louis, USA). All standards, solvents and reagents used were of highest purity (LC–MS grade where available). The water used was purified by means of a Milli-Q system (Merck Millipore, Germany).

#### Preparation of standard solutions

Stock solutions were prepared individually for each standard in a final concentration of 10 mM: 3-Haa, and Kyna were dissolved in H2O/MeOH/FA/AA (50/50/0.1/0.02). Kyn and 3-Hk were dissolved in MeOH with 0.1% FA and 0.02% AA. A final standard master mix (1 mM of each analyte) was performed by mixing individual standard stock solutions with acidified mobile phase (0.2% FA/0.05% TFA/1% MeCN in H2O). Internal standard (IS) individual stock solution (TRP_Cl) was prepared at the concentration of 84 nM in H2O/MeOH/FA/AA (50/50/0.1/0.02). All standard stocks were prepared on ice and stored at −80°C ^99^.

#### Preparation of calibration curves

Calibration curves were obtained by spiking aliquots of 50 μl of matrices (dorsal root ganglia or spinal cord samples) with 10 μl of the standard master mix solution and 10 μ of IS. The linearity of the proposed method was checked over the concentration range of 1–1700 nM to all compounds. Calibration curves were found to be linear (>0.990) over the selected range.

#### Sample preparation

Lumbar dorsal horn of the spinal cord was accurately weighted and transferred to a 2000 µl eppendorf. The samples were processed as follow: 50 μl of acidified mobile phase and 10 μl of IS were added. To each of those mixtures 150 μl of ice-cold MeOH was added and subsequently homogenized using a TissueLyser II (Qiagen). To support protein precipitation, samples were allowed to rest in −20°C for 30 min. After centrifugation (20000 g, 4°C, 15 min) supernatants were removed and evaporated to dryness under a gentle stream of nitrogen and dried extracts were reconstituted in 50 μ of acidified mobile phase. The quantification range was from 1 to 1700 nM.

#### LC–MS/MS system

The UPLC®-MS/MS analyses were performed using an Acquity UPLC (Waters, Milford, MA, USA) coupled to an Acquity TQD detector equipped with an ESI interface and a Kinetex F5 (50 × 2.1 mm, 1.7 μ mobile phase was composed of 0.2 % formic acid in water (v/v) (A) and 0.2 % formic acid in acetonitrile (v/v) (B) and it was pumped at a flow rate of 0.4 mL min-1. The column temperature was maintained at 40°C and the gradient elution program was performed as follows^99^: 0.0 min (3% B), 0.3 min (3% B), 0.8 min (30% B), 1.8 min (60% B), 2.5 min (60% B), 3.0 min (95% B), 4.4 min (95% B), 4.5 min (3% B) and 7.5 min (3% B). The samples were conditioned at 10 °C in the auto-sampler. The injection mode applied was the full-loop, using 10 μ of injection volume. Data were acquired by MassLynx v4.1 software and processed for quantification by QuanLynx V4.1 (Waters). The following generic source conditions were used in positive ionization mode: capillary voltage, 2.5 kV; cone, 10 V; desolvation temperature, 300°C; source temperature, 150°C, desolvation gas flow (N2), 750 L/h; cone gas flow, 75 L/h; collision gas (argon), 0.16 mL/min. Multiple reaction monitoring (MRM) was used for the quantification of each compound, the specific metabolite transitions are presented in **Supplementary Table 1**. The MS conditions for each analyte were determined via direct infusion of individual standard.

### Reagents

The following drugs were used in this study: 1-Methyl-tryptophan (#860646), norharmane (#N6252), Ro-618048 (#SML0233), L-kynurenine (Kyn) (#K8625), 3-Hydroxy-DL-Kynurenine (3-Hk) (#H1771), 3-hydroxyanthranilic acid (3-Haa) (#H9391) and Complete Freund’s Adjuvant – CFA; MK801 (Tocris Bioscience, Bristol, United Kingdom), diphtheria toxin (Dtx, #322326, Merck-millipore-Calbiochem San Diego, CA, USA), Kappa (k)-Carrageenan) (BDH Biochemicals, Poole, England).

### Western blot analysis

Mice were terminally anesthetized at indicated times after SNI or sham surgery and ipsilateral dorsal horn of the spinal cord tissues, draining lymph nodes (inguinal and popliteal) or dorsal root ganglion (DRGs) with dorsal root leptomeninges (DRL) ipsilateral to the lesion were harvested. Part of the experiments was performed in PBS perfused mice. The samples were homogenized, and the expression of IDO1 and KMO were evaluated using western blotting analyses. Primary culture astrocytes or cultured U87 cells were also used for western blotting analyses. Briefly, samples were homogenized in a lysis buffer containing a mixture of protease inhibitor (Protease Inhibitor Cocktail Tablets - Roche Diagnostics). Proteins were separated using SDS-polyacrylamide gel electrophoresis (SDS-PAGE-4-12%) and transblotted onto nitrocellulose membranes (Bio-rad, California, USA). The membranes were blocked with 5% dry milk (overnight) and incubated overnight at 4°C with a mouse monoclonal antibody against IDO-1 (# SC-365086 - E7 1.500, Santa Cruz Biotechnology, Dallas, Texas, USA), KMO (# NBP1-44263 1:1000, Novus Biologicals, Littleton, Colorado, USA), and a mouse monoclonal antibody against human KMO (# 60029-1-Ig 1:1000, Thermo Fisher Scientific, Massachusetts, EUA). The membranes were washed and incubated for 1 h at room temperature with an HRP-conjugated secondary antibody (1:10000; Jackson ImmunoResearch, PA, USA). Immunodetection was performed using an enhanced chemiluminescence light-detecting kit (Amersham Pharmacia, Biotech, Little Chalfont, UK) for 2 min. A mouse monoclonal antibody against β-actin (# A5316 1:10000; Sigma Aldrich, Saint Louis, Missouri, USA) and a rabbit monoclonal antibody against GAPDH (# 97166 1:5000, Cell Signaling Technology, Massachusetts, EUA) were used for loading controls. Images were used as representative blots. The capture of images was performed with Quemi TM-Doc XRS apparatus. Densitometric data were measured following normalization to the control (house-keeping gene) using Scientific Imaging Systems (Image labTM 3.0 software, Biorad Laboratories, Hercules CA).

### Culture of human astrocytes obtained from U87MG cells differentiation

U87MG cells, a human malignant glioblastoma (astrocytoma), was placed 1 x 10^5^/wells in 24 wells plate with Dulbecco’s Modification of Eagle’s Medium – DMEM (Corning Incorporated, Nova York, EUA) and 10% inactivated fetal bovine serum plus 1% Penicillin/Streptomycin. After placed, the cells were treated with 1 M of all-trans retinoic acid-ATRA (Sigma Chemical, St. Louis, MO, USA) for 7 days for astrocyte differentiation as previous described^41^. At the end, FACS analyses revealed that 98% of cultured cells express GFAP. After differentiation, cells were stimulated with human TNF recombinant (10 ng/ml) (Sigma-Aldrich Corporation, Missouri, EUA) for 24 hours.

### Construction of pLenti-GFAP-shKMO

The lentiviral vector used for cloning of shRNA targeting kynurenine 3-monooxygenase (KMO) was derived from the pLenti-GFAP-shAct1 plasmid, kindly provided by Dr Guang-Xian Zhang^42^. For pLenti-GFAP-shKMO construction, the shACT1 was replaced by the 21nt sequence GCACTGAATGCCTGCTTTCTT shRNA targeting murine KMO (NM_133809.1), designed using siRNA Wizard v3.1™ (Invivogen, San Diego, USA). The entire region of siRNA was synthesized as oligonucleotides, annealed and inserted in the previously XhoI/EcoRI digested plasmid, under standard procedures. The constructed plasmid sequence was confirmed by DNA Sanger sequencing. The vector without insertion of shKMO was used as a negative control. Oligos used for shKMO construction are listed below.

Forward:

TCGAGAAGGTATATTGCTGTTGACAGTGAGCGAGCACTGAATGCCTGC TTTCTTTAGTGAAGCCACAGATGTAAGAAAGCAGGCATTCAGTGCTGCCT ACTGCCTCGG

Reverse:

AATTCCGAGGCAGTAGGCAGCACTGAATGCCTGCTTTCTTTACATCTG TGGCTTCACTAAAGAAAGCAGGCATTCAGTGCTCGCTCACTGTCAACAGC AATATACCTTC

### Virus packaging, concentrating and tittering

The pLenti-shKMO (or empty vector) and two helper plasmids pPAX2 (Addgene #12259) and pMD2 (Addgene #12260), were used to transfect into HEK293FT cells using Lipofectamine 3000 (Thermofisher). After 6 hours of incubation, the medium with plasmids was replaced by fresh Dulbecco’s modified Eagle’s medium supplemented with 10% fetal bovine serum, 2mmol/l L-Glutamine, 100 IU/ml penicillin, and 100μg/ml Streptomycin. Supernatants were harvested after 30 hours and filtered through a 0.45µm membrane filter. The filtered supernatant was submitted to ultracentrifugation (29000 RPM, 2h, 4C). The pellet containing lentivirus was resuspended in PBS/BSA1%, and aliquots were stored at −80°C. Viral titers were assayed by Lenti-X qRT-PCR (Clontech) and titers were adjusted to 1×108 copies/ml before injection.

### Intraspinal injection of Lentivirus vector

Intraspinal injection of Lentivirus vector were performed as previous described with some modifications^44^. The mice were deeply anaesthetized by intraperitoneal (i.p.) injection of ketamine (100 mg kg−1) and xylazine (10 mg kg−1), after the mice was immobilized and attached in the rostral and caudal sites of the vertebral colum, and the skin was incised at Th12–L3. Paraspinal muscles around the left side of the interspace between Th13 and L1 vertebrae were spread and removed, the dura mater and the arachnoid membrane were carefully incised using the tip of a 30G needle to make a small cavity to allow the microcapillary Femtotip (Eppendorf, NY, USA) insert directly into the dorsal region of spinal cord. The microcapillary was inserted with 1×10^7^ lentivirus solution through the small cavity. After microinjection, the skin was sutured with 4-0 silk, and mice were kept on a heating pad until recovery.

### Intrathecal ShRNA against KMO

The short hairpin RNA targeting the murine expression of KMO (NM_133809.1) was designed and acquired from Genecopoeia (plasmid reference MSH037508-CU6). The *in vivo* transfection was performed using InVivo JetPei reagent (Polyplus) and administered intrathecally as described previously^100^ starting at 10 up to 13 days after SNI induction.

### Real-time RT-PCR

After collection of tissue from the region comprised between lumbar segments (L3-L6) of the dorsal horn of spinal cord, draining lymph nodes ipsilateral to the lesion, primary culture astrocytes or cultured U87 cells were rapidly homogenized in Trizol (Sigma) reagent at 4°C. Then, total cellular RNA was purified from tissue, according to the manufacturer’s instruction. RNA concentration was determined by optical density at a wavelength of 260 nm by means of the apparatus NanoVue ® (*GE Helthcare*). One microgram of total RNA was transcribed to cDNA by reverse transcriptase enzyme action Improm Pre-II ® (Promega, Madison, Wisconsin, USA). Quantitative RT-PCR reaction in real time was done on an ABI Prism ® 7500 Sequence Detection System (*Applied Biosystems*), using System SYBR-green fluorescence (*Applied Biosystems, Warrington, UK*) for the quantification of amplification. RT-PCR was performed with the final volume of the reaction 6.25 μL and kept on 95 °C (10 min) and 40 cycles of 94 °C (1 min), 56 °C (1 min) and 72 °C (2 min). The melting curve was analyzed (65-95 °C) to verify that only one product was amplified. Samples with more than one peak were excluded. The results were analyzed by the method of quantitative relative expression 2^-ΔΔCt^ as previously described^101^. The primer pair for mouse can be found at **Supplementary Table 2.**

### Immunofluorescence

After appropriate times, animals were deeply anesthetized with ketamine and xylazine and perfused through the ascending aorta with PBS, followed by 4% paraformaldehyde. Spinal cord sections (60 μ (20 μm) were washed in PBS (0.01 M, pH 7.4) 3 x 5 minutes and incubated in 1% BSA, dissolved in phosphate buffered saline with Triton X100 (PBST) at 0.1% for 1 hour. In some experiments related to DRLs, whole lumbar vertebrae containing spinal cord and DRGs (L3-L5) referring to sciatic nerve were obtained from SNI (14 days) and sham animals were fixed with 4% paraformaldehyde buffered with 0.1M phosphate solution, pH 7.2-7.4, for 4 hours at 4°C. Next, tissues were decalcified with 5% EDTA in 1X Dulbecco’s PBS solution for 10 days at 4°C and serially cryoprotected in 10%, 20%, and 30% sucrose serie diluted in 1X Dulbecco’s PBS solution overnight each at 4°C. Subsequently, the sections were washed in PBS (0.01 M, pH 7 4) 3 x 5 minutes and then processed according to the technique of immunofluorescence labeling with overnight incubation at 4 °C with the polyclonal anti-Iba1 1:250 (# NB100-1028, Novus Biologicals, Littleton, Colorado, USA), monoclonal anti-GFAP 1:500 (# MAB3402X, Millipore, Darmstadt, Germany), anti-NeuN 1:200 (# MAB377X Millipore, Darmstadt, Germany), anti-KMO 1:200 (# NBP1-44263, Novus Biologicals, Littleton, Colorado, USA), a mouse monoclonal antibody against IDO-1 (# sc365086, clone E7, Santa Cruz Biotechnology, Texas, EUA) and polyclonal anti-GFP conjugated Alexa 488 (# A211311, Life Technologies, California, EUA) and CD11c 1:300 (#117308, Biolegend, California, EUA). After incubation with the primary antibodies, the sections were washed in PBST 3 x 5 min and incubated at room temperature for 1 hour with Alexa fluor 488^®^goat anti-mouse or 594^®^donkey anti-rabbit IgG (Molecular Probes, Eugene, Óregon, USA). The sections were washed with PBST as described earlier, and mounted on glass slides, covered with cover slips with FluromountTM Aqueous Mounting Medium (Sigma, St. Louis, Missouri, USA). Sections of spinal cord were analyzed by confocal microscopy (SP5, Leica, Wetzlar, Germany).

### Generation of bone marrow-chimeric mice

Recipient mice were exposed to 9-gray total-body irradiation using a X-ray source (Mark I, model 25). One day later, the animals were injected via tail vein with 4 × 106 bone marrow (BM) cells freshly collected from donor mice. The cells were aseptically harvested by flushing femurs with Dulbecco’s PBS (DPBS) containing 2% fetal bovine serum. The samples were combined, filtered through a 40 μm nylon mesh, centrifuged, and passed through a 25-gauge needle. Recovered cells were resuspended in DPBS at a concentration of 5 × 106 vial nucleated cells per 200 μl. Irradiated mice transplanted with this suspension were housed in autoclaved cages and treated with antibiotics (10 mg of ciprofloxacin per milliliter of drinking water given for 2 weeks after irradiation)^100^. Mice were subjected to SNI surgery 2 months after transplantation. The chimeric mice were generated as follow (donor → recipient): (1) WT and Ido1^-/-^ recipient mice transplanted with BM from WT mice (WT → Ido1^-/-^ and WT → WT), (2) Ido1^-/-^ recipient mice transplanted with Ido1^-/-^ donor mice (Ido1^-/-^ →Ido1). At 60 days after transplantation, SNI was induced in chimeric mice.

### Generation of CD11c^DTR^*^/^*^hema^ chimeric mice and dendritic cells depletion

For *in vivo* depletion of DCs we performed a previously described method^31, 102^, with modifications. We used a chimera-based approach to avoid side effects during DCs depletion protocol. The experimental design for the generation of chimeric mice is shown in Fig. 2D. Briefly, WT mice were irradiated with a dose of 9.0 Gy. After 24 h, a total of 5.0 x 10^6^ bone marrow-derived cells from CD11c-DTR mice were injected intravenously in the irradiated mice. After 8 weeks, the reconstituted chimeric mice (CD11c^DTR/hema^ mice) were randomly separated into for groups and submitted to following experimental protocols. For conditional DC ablation [CD11c^DTR/hema^ mice chimeras were inoculated i.p. with 16 ng DTx/g body weight every second day from day 0 up to day 12 after SNI induction.

### Dendritic cells sorting from draining lymph nodes

At 14 days after SNI, draining lymph nodes (DLNs; inguinal and popliteal) from CD11c-eYFP mice were homogenized in RPMI 1640 media pH 7.4 (GIBCO), then digested with 1 mg/ml of collagenase A for 30 min at 37°C. Single cell suspensions were prepared using a 70 μm-Cell strainer (Corning– Durhan, USA) nylon mesh, layered over RPMI 1640 containing 10% FCS and centrifuged at 450 *g* for 10 min at 4°C and the supernatant was discarded. To purify total CD11c^+^ DCs, single cell suspensions from lymph nodes were further sorted (eYFP^+^ cells) in a FACSAria III sorter. Over 5% of the sorted cells obtained were CD11c^+^. Sorted cells were submitted to RNA extraction, reverse-transcribed with High Capacity Kit (Life Techonologies) and analyzed by quantitative RT-PCR with a Step One Real-time PCR system as described above (Applied Biosystems).

### Microglia and astrocytes cells sorting from spinal cord

Astrocytes and microglia were isolated from the spinal cord as previously described^40, 43, 103^; with minor modifications. Briefly, the spinal cord between lumbar segments (L4-L6, pooled of 8 animals) were collected from CX3CR1^GFP/+^ mice and incubated isolated in 1 ml of RPMI medium containing 1mg/ml of collagenase D (Roche, cat. no. 1-088-874) for 30 minutes at 37°C. After this time, the tissues were passed through a cell strainer (100 µm), followed by centrifugation with RPMI medium containing 10% fetal bovine serum (FBS, Gibco). Cells were obtained and resuspended in 10 ml of 33% Percoll (GE Health Care) solution. After centrifugation, cells were resuspended and incubated in PBS 0.1M (2% FBS) containing cell viability dye – APCH7 (cat. no. 65-0865-14, eBioscience, 1:5000) and anti-ACSA2 - PE (IH3-18A3, Miltenyi Biotec, 1:250), a specific marker of astrocytes, for 15 minutes at 4°C. Finally, cells were isolated using FACSAria III (BD Bioscience) and then the pellets obtained were resuspended in lysis buffer to isolate mRNA. The data were analyzed using FlowJo 10 software (Treestar, Ashland, USA). The gating strategy used to perform cell sorting from spinal cord is depicted in Fig. 7. Expression of specific cell markers, *Cx3cr1* for microglia and *Gfap* for astrocytes and *Kmo* were analyzed by real-time PCR (see specific method) in both populations.

### Flow cytometry acquisition

DLNs (inguinal and popliteal) were removed from CD11c^DTR/hema^ mice treated with Dtx or vehicle after 14 days of SNI or sham surgery, teased into single-cell suspensions, and filtered through a 70-μm cell strainer (Corning # 431751 – Durhan, USA) and proceed as described above. DCs (CD11c-eGFP^+^) cells were detected and the frequency of cell group was determined in BD FACSVerse (BD Biosciences), and data were analyzed using FCS Express (De Novo Software™).

### Generation of bone marrow-derived dendritic cells (BMDCs) and transfer

BMDCs were generated from the differentiation of bone marrow cells. Briefly, animals were killed and the femur bone carefully dissected. In sterile environment, the bone epiphyses were cut and a 20g needle, coupled to a complete RPMI-1640 filled syringe (supplemented with 10% fetal bovine serum, glutamine (2 mM), streptomycin-0.01 mg/mL, penicillin-10 U / mL, antifungal-amphotericin B-30 μg / mL) was inserted at one end. Next, the internal cavity of the bone was washed away, removing the entire bone marrow. Cells were added to the culture plates at the density of 2.0 x 10^6^ cells / dish in a final volume of 10 mL of complete RPMI-1640 supplemented with 20 ng/mL GM-CSF (granulocyte-macrophage colony stimulating factor; R&D Systems). After 3 days, another 10 mL of medium supplemented with the same concentration of GM-CSF was added. At the end of 7 days, the cells were collected from the plate and used for the remaining experiments^31^. BMDCs (5.0 x 10^6^; i.v.) were transferred to WT or *Ido1^-/-^* mice one day after SNI surgery.

### Primary Astrocytes Culture and Microglia Conditional Medium (MCM)

For the primary culture, the astrocytes were obtained by the microbeads isolation on the cortex of newborn mice (1 to 3 days). The newborn mice were anesthesia in ice, and the cortex was collected in Dulbecco’s Modification of Eagle’s Medium – DMEM (Corning Incorporated, Nova York, EUA) with penicillin and without bovine fetal serum. After isolation, the cells of cortex were dissociated and marker with anti-GLAST (ACSA-1) biotinilad antibody, after with anti-biotin microbeads (Miltenyi Biotec, Bergisch Gladbach, Germany). The cells were passed in MS Columns (Miltenyi Biotec, Bergisch Gladbach, Germany), and only the ACSA-1 positives cells stay attached in the columns. The isolated astrocytes were placed 5×10^4^ cells/well in a 24 well place for 7 days with DMEM (Corning Incorporated, Nova York, EUA) (high glucose, 10% heat-inactivated fetal bovine serum and 1% Penicillin/Streptomycin). After 7 days, the astrocytes were stimulated with TNF recombinant (10 ng /ml) (Sigma-Aldrich Corporation, Missouri, EUA) or MCM (300 µl) for 24hours. To determinate the purity of culture, the cells were marker with anti-GFAP-PE (# 561483, California, EUA, BD Biosciences) for flow cytometry acquisition. For the MCM, microglia were obtained by the same protocol of astrocytes primary culture. However, the cells of newborn mice cortex were dissociated and marker with anti-CD11b-PE antibody, after with anti-PE microbeads (Miltenyi Biotec, Bergisch Gladbach, Germany). The cells were passed in MS Columns (Miltenyi Biotec, Bergisch Gladbach, Germany), and only the CD11c positives cells stay attached in the columns. The isolated microglia were placed 10^5^ cells/well in a 24 wells place for 5 days with DMEM (Corning Incorporated, Nova York, EUA) (high glucose + 10% heat-inactivated fetal bovine serum + 1% Penicillin/Streptomycin). After 5 days, the microglia were stimulated with LPS (100 ng /ml) (Sigma-Aldrich Corporation, Missouri, EUA) for 30 min, washed and stayed in medium for 24 hours to obtain the MCM.

### Data analyses and statistics

Data are reported as the means ± s.e.m. The normal distribution of data was analyzed by D’Agostino and Pearson test. Two-way analysis of variance (ANOVA) was used to compare the groups and doses at the different times (curves) when the responses (nociception) were measured after surgery or treatments. The analysed factors were the treatments, the time, and the time versus treatment interaction. If there was a significant time versus treatment interaction, One-way ANOVA followed by Bonferroni’s t-test was performed for each time. Alternatively, if the responses (eg. protein expression, mRNA expression) were measured only once after the stimulus injection, the differences between responses were evaluated by one-way ANOVA followed by Bonferroni’s t-test (for three or more groups), comparing all pairs of columns or when appropriate by two-tailed unpaired Student’s t-test. For electrophysiology data, the paired sample t-test was used. P values less than 0.05 were considered significant. Statistical analysis was performed with GraphPad Prism (GraphPad Software, San Diego, CA, USA). No statistical methods were used to predetermine sample size. Variation within each data set obtained by experiments with mice or primary cells was assumed to be similar between genotypes since all strains were generated and maintained on the same pure inbred background (C57BL/6).

## Data availability

The data supporting the findings of this study are available within the article and its supplementary information files or from the corresponding authors on reasonable request.

## Acknowledgments

The authors gratefully acknowledge the technical assistance of Ieda R. Schivo, Sergio R. Rosa and Eleni Tamburus. The research leading to these results received funding from the São Paulo Research Foundation (FAPESP) under grant agreements n° 2011/19670-0 and 2014/50265-3 (Thematic Projects) and 2013/08216-2 (Center for Research in Inflammatory Disease) and a CNPq grant n° 304883/2017-4. ALM is supported by grants from the NIH (AI103347) and CRUK (CIP award).

## Author contributions

G.R.S, A.G.M, M.D.F and A.D designed, performed experimental work, analyzed data and prepare the manuscript. R.M.G and M.D.F performed experimental work related to FACS sorting and PCR and analyzed data.. A.H.L, F.I.G, R.L.S, D.A.S, J.T, G.P, D.S.C, M.B.S, A.S.M, and R.K performed experiments. L.M.M, G.A.B and N.P.L performed experiments related to mass spectrometry and analyze the data L.M.A and N.T.C performed experiments related to lentivirus and ShRNA. H.L and L.H. performed experiments and important scientific comments. J.C.A.F., F.Q.C., R.L, and J.C.O.C provided critical materials and comments. A.M designed and supervised the study and provided critical materials and comments. T.M.C designed, directed and supervised the study, interpreted data and wrote the manuscript. All authors reviewed the manuscript and provide final approval for submission.

