## Supplementary material for "Meningeal dendritic cells-astrocytes interactions elevate the kynurenine metabolic pathway to sustain neuropathic pain": Suplemmentary figures and tables (from another version of this preprint: version 1, 2021-01-26)

**This PDF file includes:**

Materials and Methods

Figs. S1 to S13

Captions for Movies S1 to S3

**Other Supplementary Materials for this manuscript include the following:**

Movies S1 to S3

**Supplementary Figures**

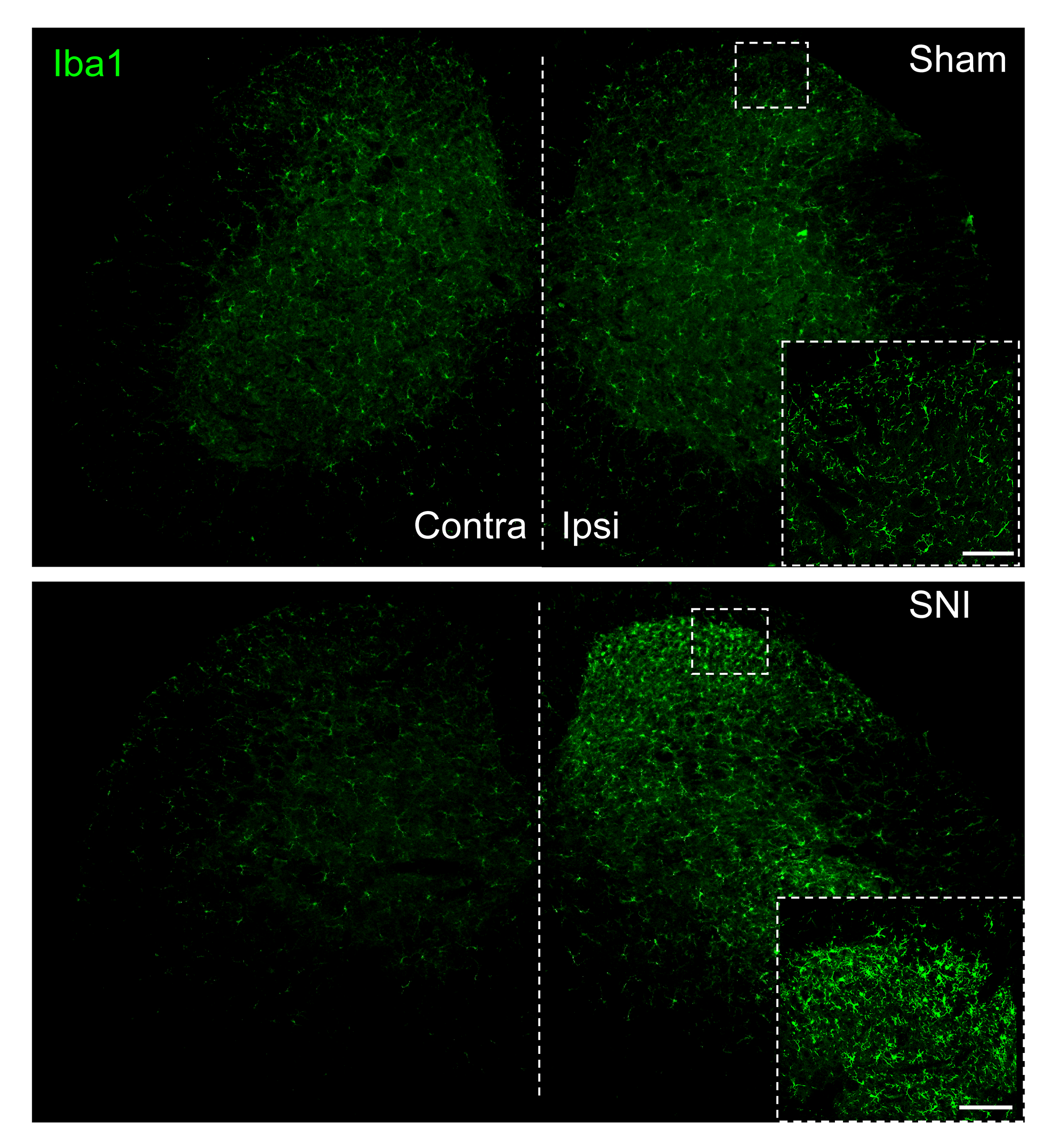

**Supplementary Fig. 1.** Microglial cells activation/proliferation in the spinal cord after spared nerve injury (SNI). Representative confocal images of the spinal cord at 10 days after SNI in WT mice. IBA1+ microglia are shown in green. Dotted boxes show regions of higher magnification in the dorsal horn (right panels). Scale bars: 250 µm (left panels) and 50 µm (right panels).

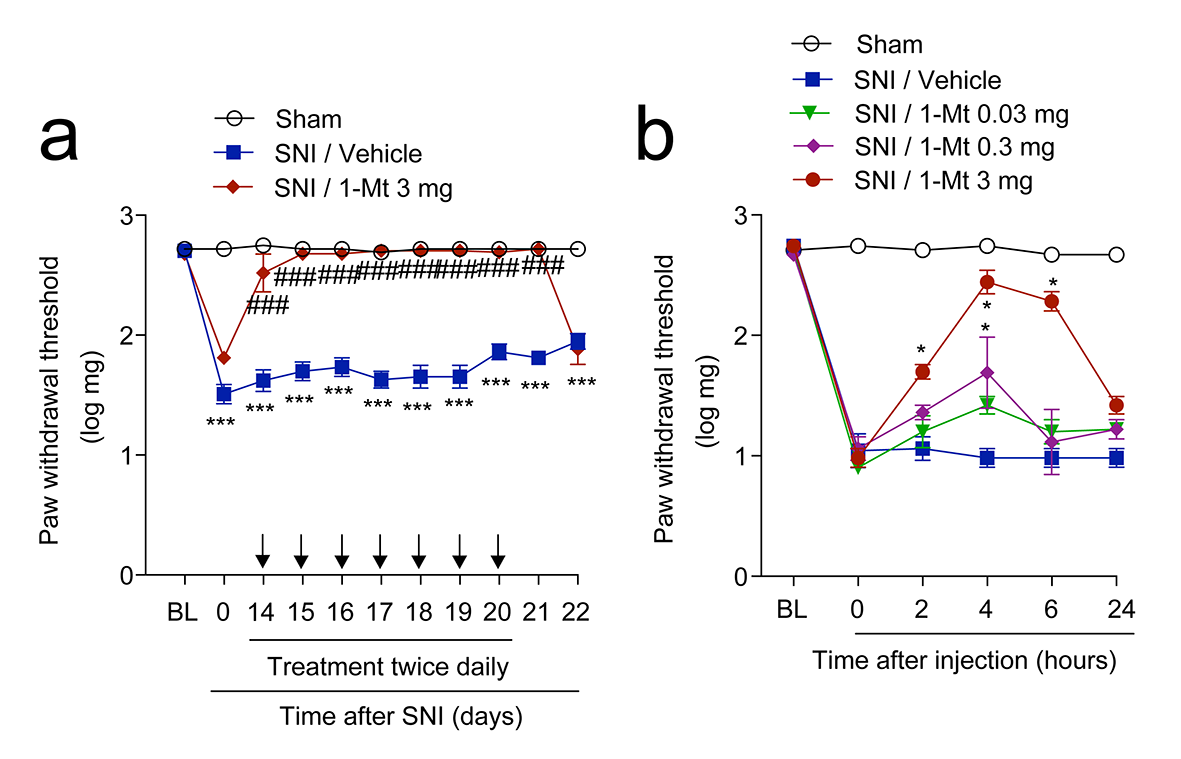

**Supplementary Fig. 2. (a)** Mechanical nociceptive threshold was determined before and 14 days after SNI. Mice were treated (i.p.) with vehicle or 1-Mt 3 mg twice daily during seven consecutive days and mechanical allodynia was measured up to 22 days after SNI (n=6). **(b)** Mechanical nociceptive threshold was determined before and 21 days after SNI. Mice were treated i.p. with vehicle or 1-methil-DL-tryptophan and mechanical allodynia was measured up to 24 h after treatment (1-Mt, 3 mg/per mouse) (n=6). Data are expressed as the mean ± s.e.m. **P* < 0.05, ****P* < 0.001 versus sham ###*P* < 0.001 versus vehicle treated. Two-way ANOVA, Bonferroni’s post-test (a and b).

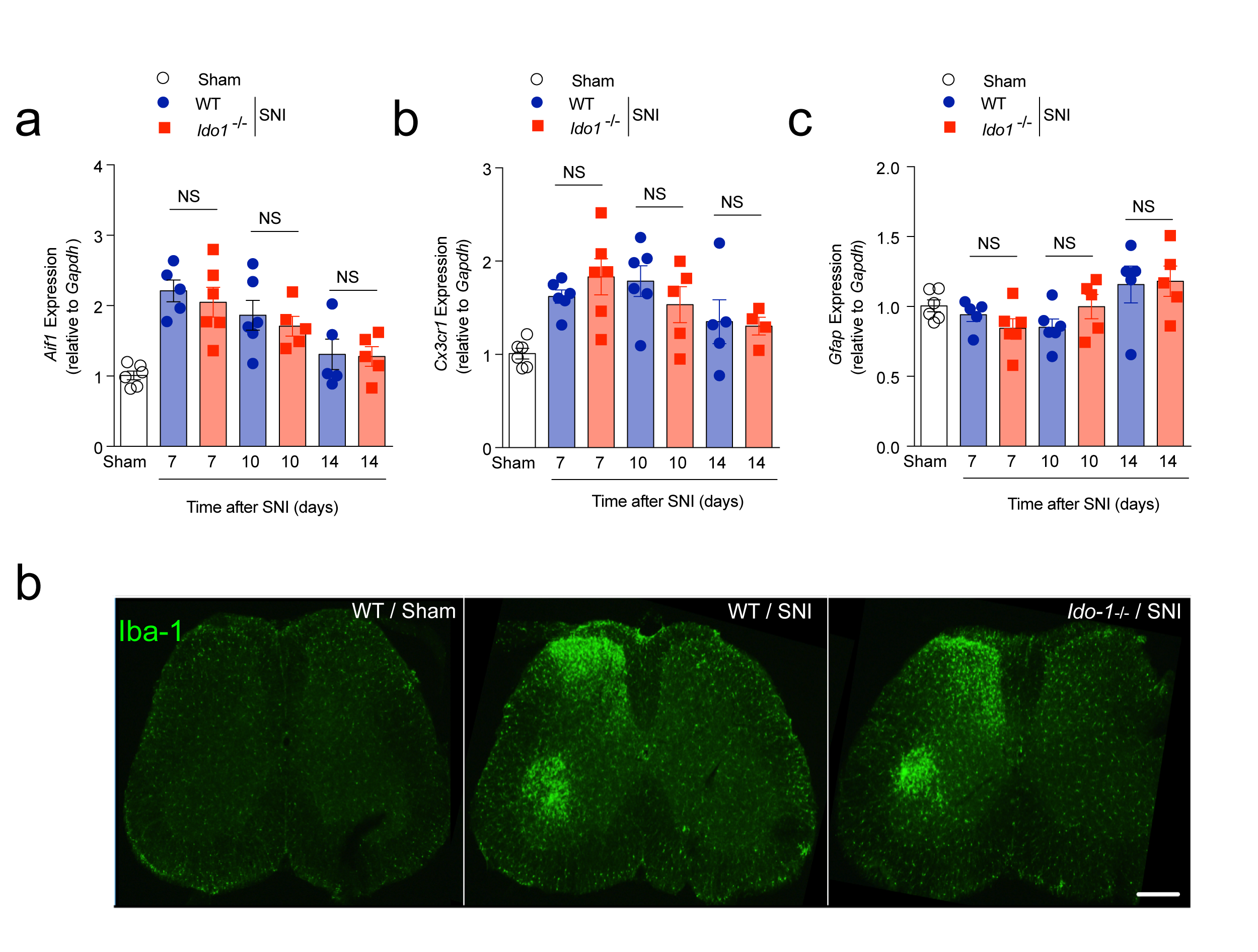

**Supplementary Fig. 3. Kynurenine metabolic pathway did not participate in the activation of glial cells in the spinal cord after SNI.** **(a-c)** The *Aif1, Cx3cr1 and Gfap* mRNA expression in the ipsilateral dorsal horn of the spinal cord (7, 10 and 14 days) after sham or SNI surgeries in WT or *Ido1*^-/-^ mice (n=5-6) **(d)** Representative images showing immunoreactivity for IBA-1 (green color) on spinal cord sections from WT and *Ido1*^-/-^ mice at 10 d after SNI or sham-operated mice (n=3). Scale bars: 300 μm NS = non-significative, One-way ANOVA, Bonferroni’s post-test (a-c; means ± s.e.m.).

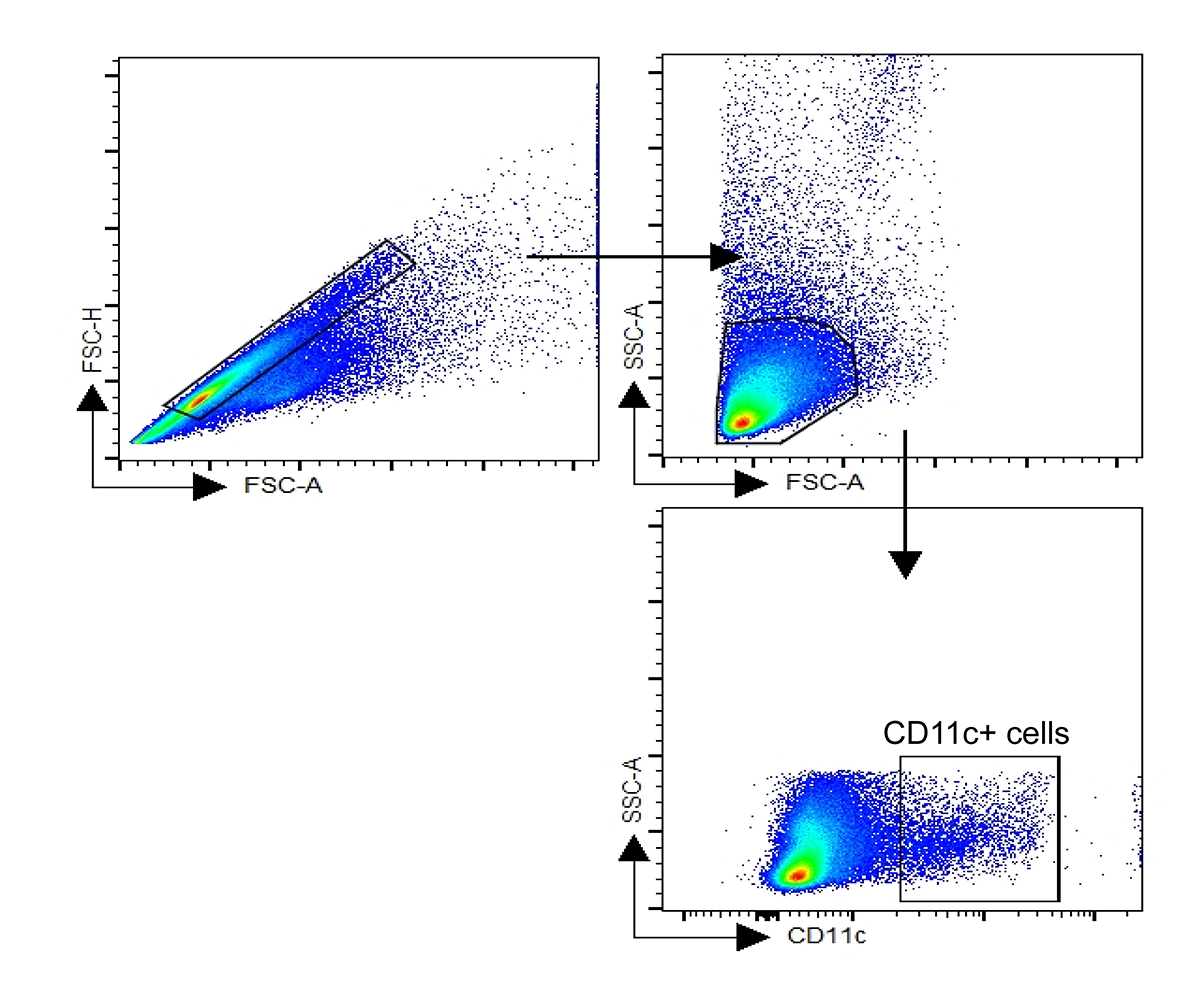

**Supplementary Fig. 4.** Representative FACS sorting strategy for CD11c^+^ cells isolation from draining lymph nodes (popliteal and inguinal).

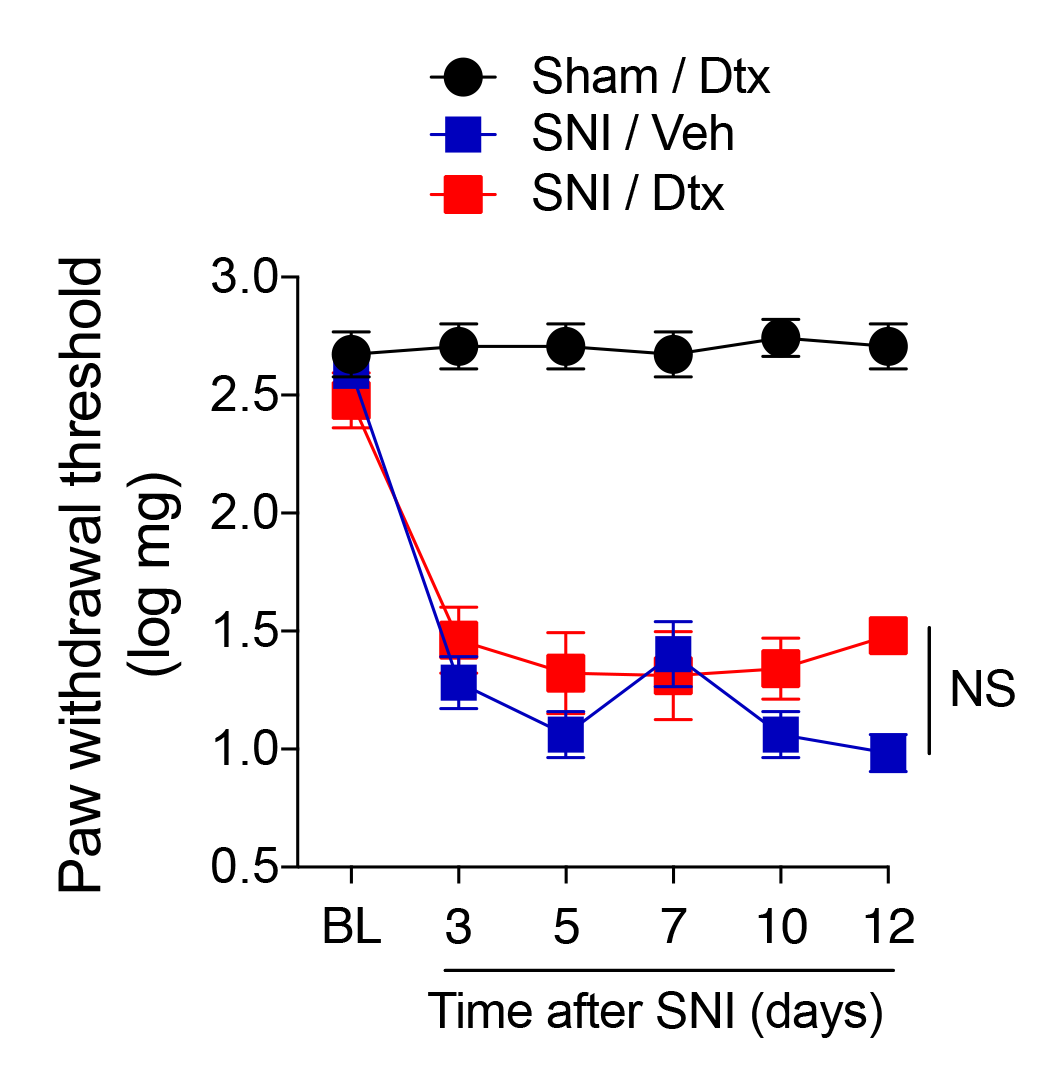

**Supplementary Fig. 5**. **Diphtheria toxin treatment did not change mechanical allodynia in WT mice.** Mechanical nociceptive threshold was determined before and up 21 days after SNI. One group of mice was treated with 16 ng Dtx/g body weight (i.p.) every second day from day 0 up to day 12 after SNI induction (n=6). NS = non-significative; Two-way ANOVA, Bonferroni’s post-test (means ± s.e.m.).

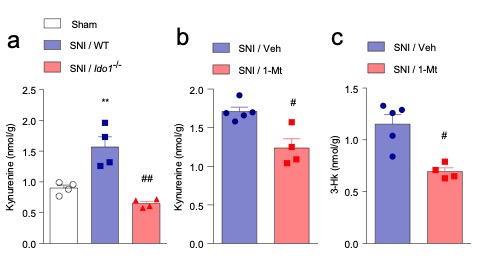
 **Supplementary Fig. 6. Kyn and 3-Hk levels in the spinal cord after SNI: role of IDO-1 expressed in DRLs.** Levels of kynurenine (Kyn) was determined in the ipsilateral dorsal horn of the spinal cord of **(a)** WT or *Ido*1^-/-^ mice or in **(b)** WT mice treated with vehicle or 1-methil-DL-tryptophan (1-Mt, 15 µg/per site) after SNI surgeries (14 days after surgeries; n=4-5). **(c)** 3-Hydroxykynurenine (3-Hk) levels in the ipsilateral dorsal horn of the spinal cord of WT mice treated with vehicle or 1-methil-DL-tryptophan (1-Mt, 15 µg/per site) after SNI surgeries (14 days after surgeries; n=4-5) **P*< 0.05 versus sham group. # *P*< 0.05, ## *P*< 0.01 versus vehicle treated. One-way ANOVA, Bonferroni’s post-test (a; means ± s.e.m.). Unpaired Student’s *t-*test (b and c; means ± s.e.m.).

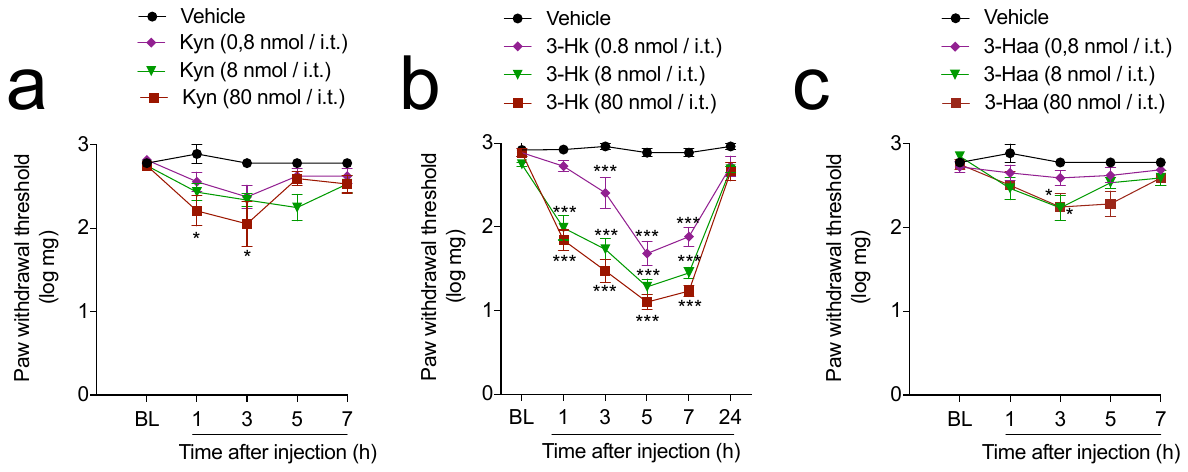

**Supplementary Fig. 7. Spinal effects of kynurenine metabolites in mechanical nociceptive threshold.** Mechanical nociceptive threshold was evaluated before and up to 07 h after intrathecal injection of equimolar doses of (a) kynurenine (Kyn, 0.8-80 nmol), (b) 3-hydroxykynurenine (3-Hk, 0.8-80 nmol); (c) 3-Hydroxyanthranilic acid (3-Haa – 0.8-80 nmol) or vehicle (saline) in WT mice (n=4-6). **P*< 0.05, ****P*< 0.001 versus vehicle treated. Two-way ANOVA, Bonferroni’s post-test (means ± s.e.m.).

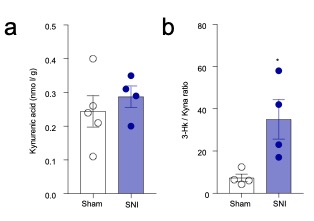

**Supplementary Fig. 8. Kyna levels and its relationship with 3-Hk in the spinal cord after SNI (a)** Levels of kynurenic acid (Kyna) were determined in the ipsilateral dorsal horn of the spinal cord of mice after sham or SNI surgeries (14 days after surgeries; n=4-5). **(b)** Ratio between 3-Hk and Kyna levels in the ipsilateral dorsal horn of the spinal cord of mice after sham or SNI surgeries (14 days after surgeries; n=4). **P*< 0.05 versus sham group. Unpaired Student’s *t-*test (means ± s.e.m.).

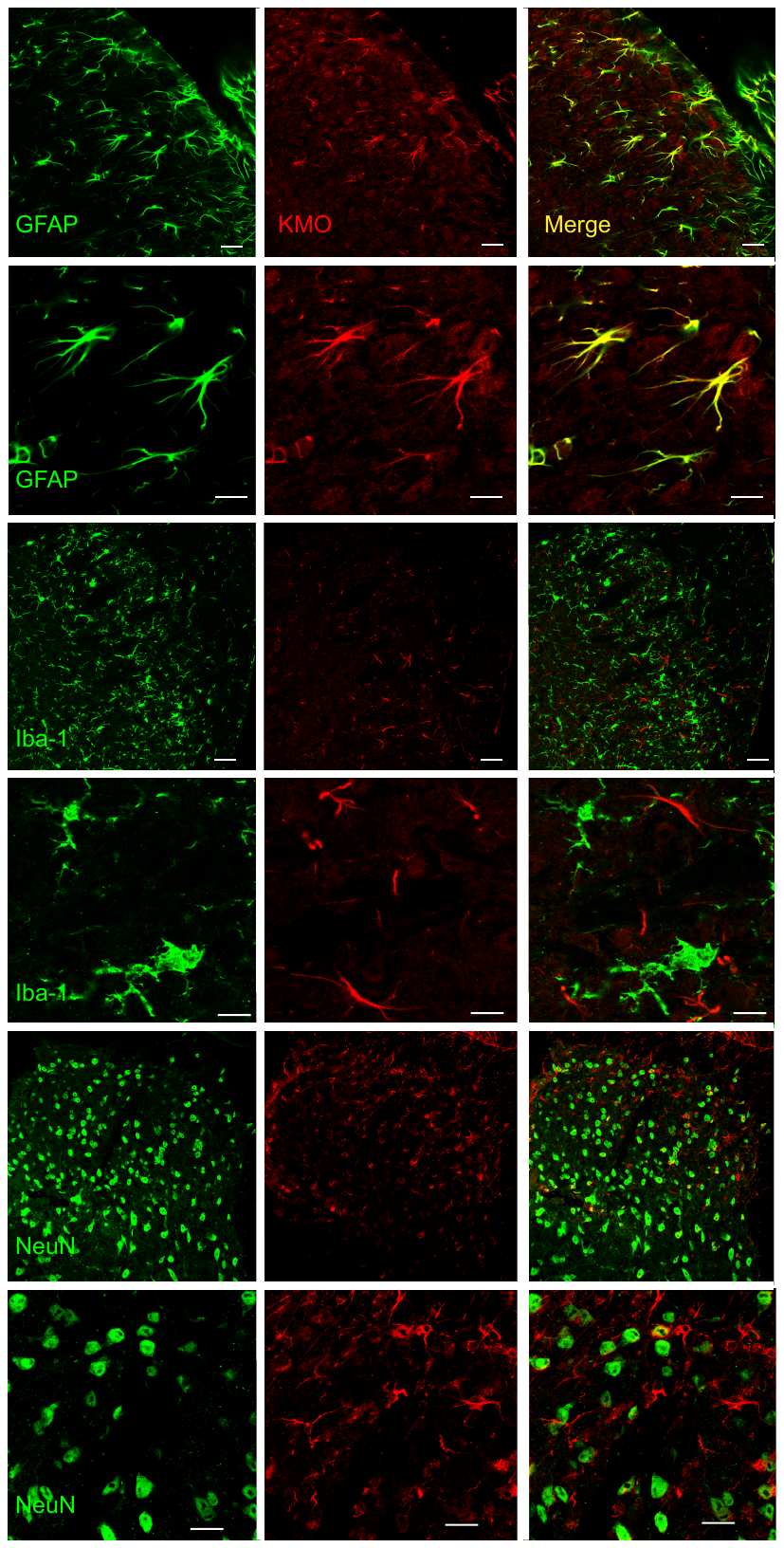

**Supplementary Fig. 9. Cellular pattern of KMO expression in the spinal cord after peripheral nerve injury.** At 14 days after sham or SNI surgeries, the expression of KMO was analyzed by immunofluorescence. Representative of 4, bar scale: 50 and 25 µm.

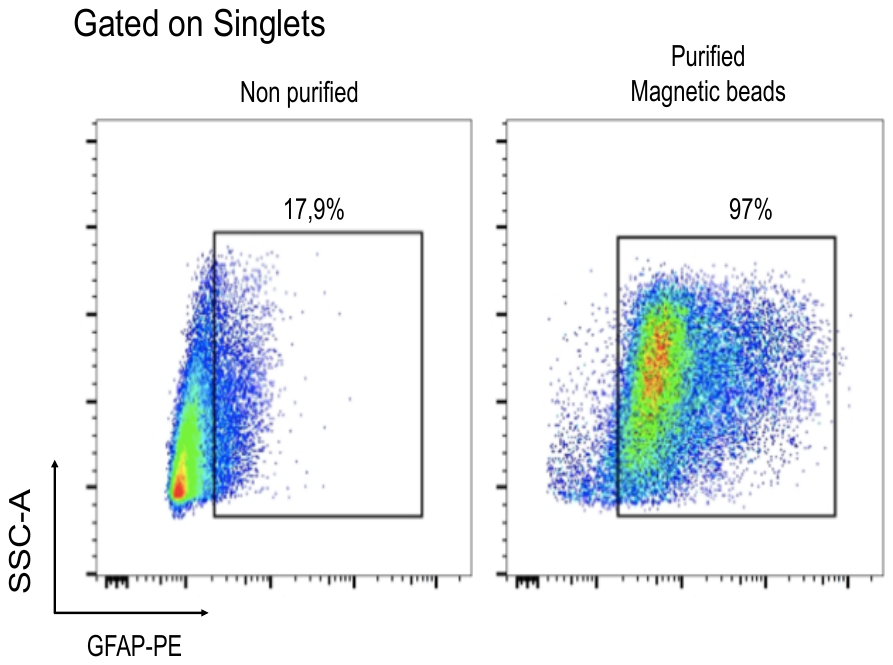

**Supplementary Fig. 10.** Representative FACS strategy for GFAP^+^ cells showing the efficacy of the purification protocol. Newborn mouse brain cortex cells were isolated and purified as described in the methods section. The FACS panels depicted the percentage of GFAP^+^ cells before and after purification.

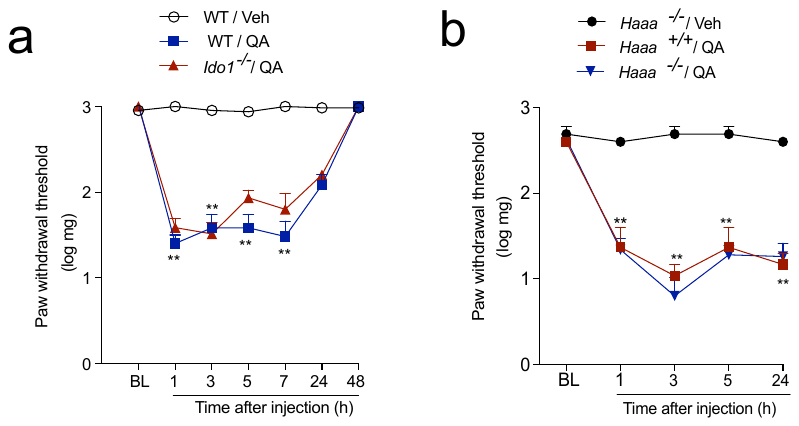
 **Supplementary Fig. 11.** Mechanical nociceptive threshold was evaluated before and after intrathecal injection of QA (80 nmol) or vehicle (saline) in WT and **(a)** *Ido1*^-/-^ mice or in **(b)** in *Haao*^+/+^ and *Haao*^-/-^ mice (n=4-5). ***P*< 0.001 versus vehicle treated. NS = non significative. Two-way ANOVA, Bonferroni’s post-test (means ± s.e.m.).

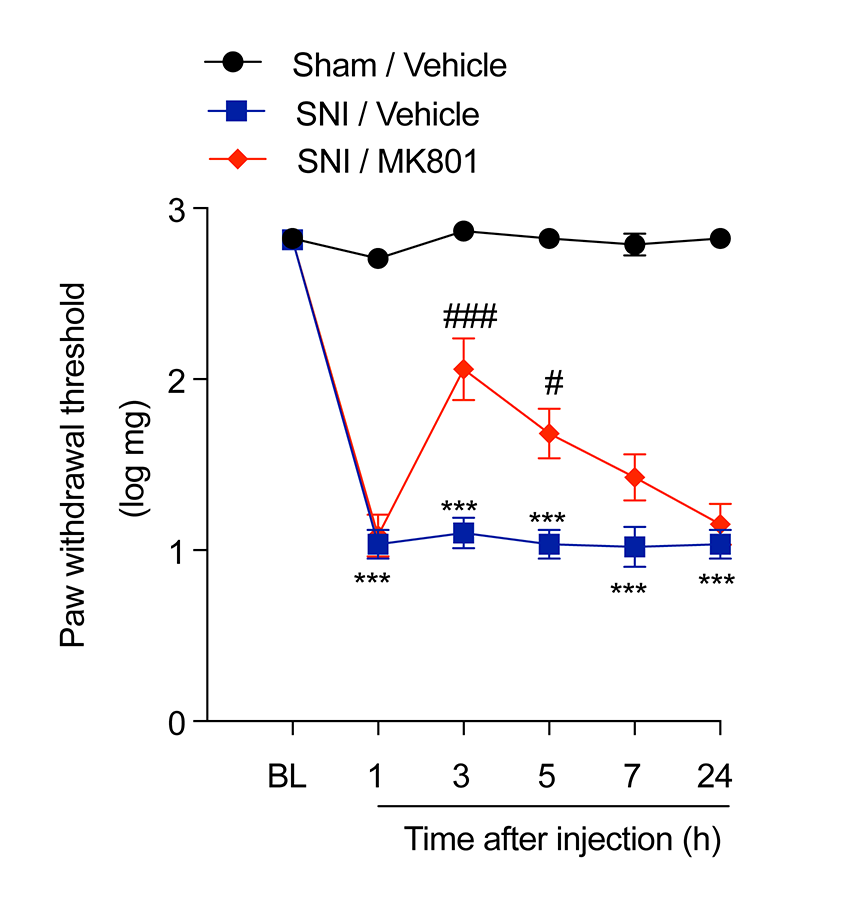

**Supplementary Fig. 12. NMDA receptors are involved in the maintenance of SNI-induced neuropathic pain.** Mechanical nociceptive threshold was determined before and 14 days after SNI. Mice were then treated intrathecally (i.t.) with MK801 (NMDA receptor antagonist, 10 nmol) or vehicle and mechanical allodynia was measured up to 24 h after treatment (n=5-6). ****P*< 0.001 versus sham group. ^#^*P*< 0.05, ^##^*P*< 0.01, SNI vehicle treated. **P*< 0.05 versus baseline. Two-way ANOVA, Bonferroni’s post-test (means ± s.e.m.).

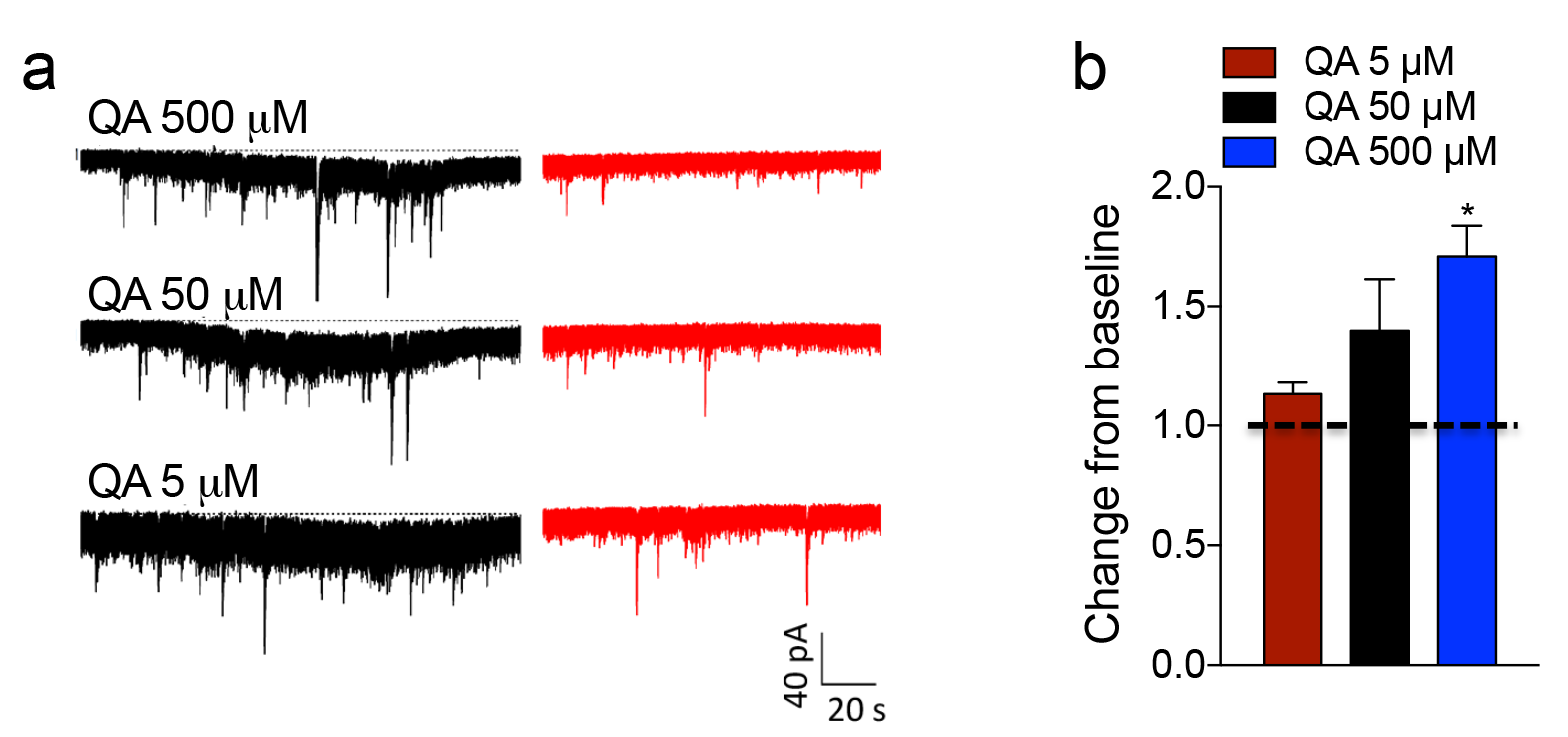

**Supplementary Fig. 13. QA elicits an inward currents in neurons from spinal cord slices** (a) Representative traces in black show the development of an inward current upon application of QA in different concentrations (5, 50 and 500 µM – black traces) and the drug washout recorded between each application (red traces) (b) Current values upon application of QA are shown on the right normalized using the control baseline as reference (n=4). **P*< 0.05 versus baseline. The paired t-test (means ± s.e.m.).

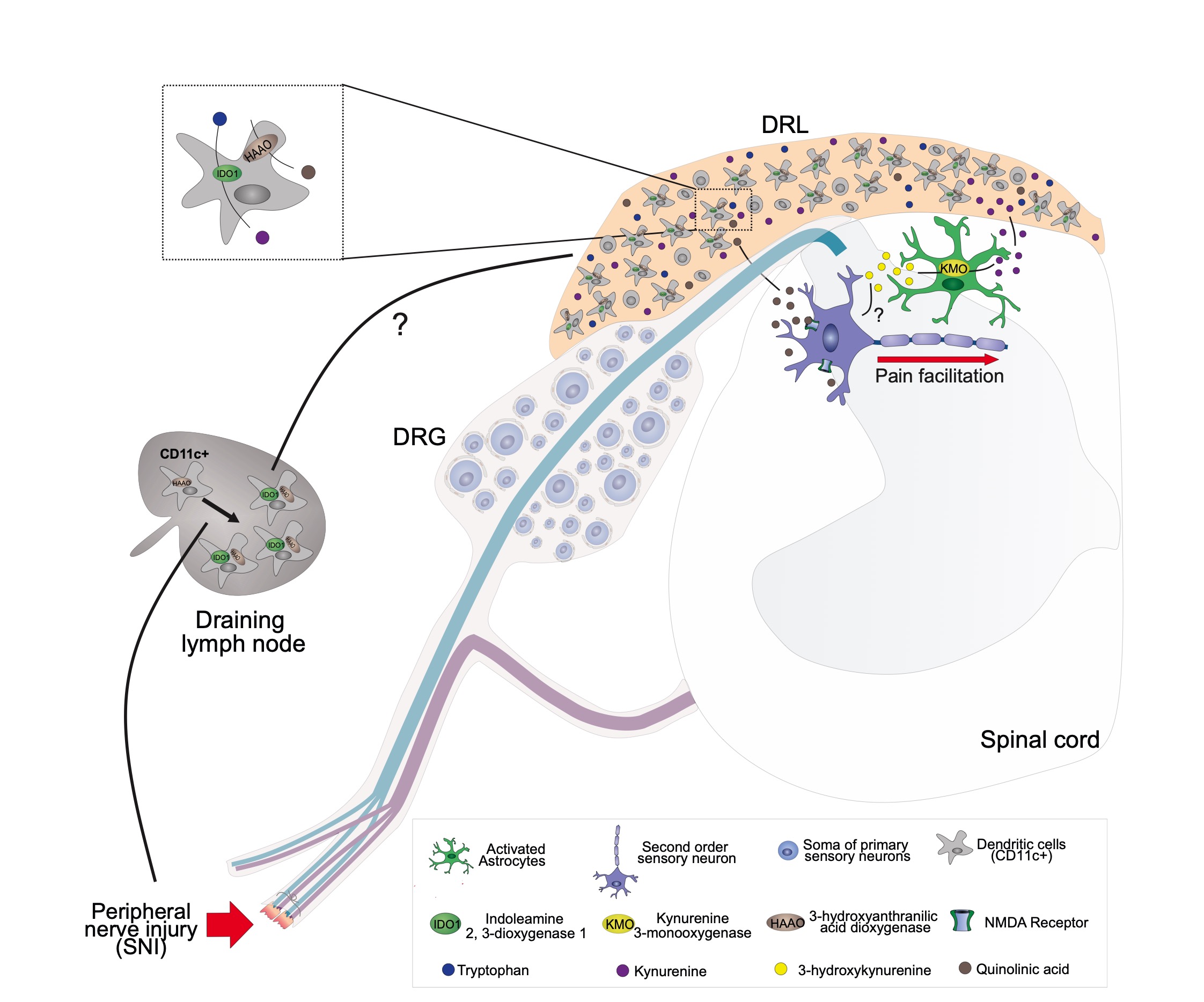

**Fig. 14. Schematic representation of the role of kynurenine pathway in the maintenance of neuropathic pain.** After peripheral nerve injury, there is an up regulation of IDO1 in dendritic cells (DCs), which accumulate in the dorsal root leptomeninges (DRLs) leading to an increase in the levels of kynurenine (Kyn) in the spinal cord. In the spinal cord, Kyn is metabolized by astrocytes-expressed KMO into a potent pronociceptive metabolite, 3-Hydroxykynurenine (3-Hk). Additionally, HAAO-expressing DCs in (DRLs) might be responsible to increase the levels of QA in the spinal cord. QA is also a pro-nociceptive molecule in the spinal cord acting through an enhancement of NMDA glutamatergic transmission and consequently to maintain pain hypersensitivity.

**Supplementary Table 1.** Compound dependent parameters in MRM mode for UPLC**^®^**- MS/MS analysis.

| Analyte | MRM (*m/z*) | Cone (V) | Collision energy (eV) |
| --- | --- | --- | --- |
| Kyn | 209 > 94  209 > 146  209 > 192 | 16 | 10 |
| 3-Hk | 225 > 110  225 > 162  225 > 208 | 5 | 15 |
| Kyna | 190 > 89  190 > 116  190 > 144 | 26 | 18 |
| 3-Haa | 154 > 80  154 > 108  154 > 136 | 14 | 12 |
| QUIN | 168 > 106  168 > 124  168 > 150 | 16 | 8 |
| TRP_Cl | 239 > 180  239 > 193  239 > 222 | 10 | 15 |

**Supplementary Table 2.** List of real-time PCR primers

|  | **Gene** | **Sequence** |
| --- | --- | --- |
| **Mouse** | *Ido1* | fwd: 5’-TGCCCGACGCATACACC- 3’ |
|  |  | rev: 5’-GTCCGTCCGTGCTCAGTGG- 3’ |
|  | *Kmo* | fwd: 5’-TGTGCCCATGAAAGCCAGAA- 3’ |
|  |  | rev: 5’-GCACCTTCGACTTGGCATAG- 3’ |
|  | *Haao* | fwd: 5’-GCAACAAGCCTTATGCACCAGG- 3’ |
|  |  | rev: 5’-ACATCCACATCCTGCCTTGG- 3’ |
|  | *Gfap* | fwd: 5’-AGGGCGAAGAAAACCGCATCACC- 3’ |
|  |  | rev: 5’-TCTAAGGGAGAGCTGGCAGGGCT- 3’ |
|  | *Cx3cr1* | fwd: 5’-GCCTCTGGTGGAGTCTGCGTG- 3’ |
|  |  | rev: 5’-CGCCCAAATAACAGGCCTCAGCA- 3’ |
|  | *Aif1* | fwd: 5’-TGAGGAGCCATGAGCAGCCAAAG- 3’ |
|  |  | rev: 5’-GCTTCAAGTTTGGACGGCAG- 3’ |
|  | *Gapdh* | fwd: 5’- CATCTTCTTGTGCAGTGCCA- 3’ |
|  |  | rev: 5’- CGGCCAAATCCGTTCAC- 3’ |

**Movies Captions**

Movie 1. KMO^+^ (red) and GFAP^+^ (green) cells in the spinal cord

Movie 2. IBA-1^+^ microglia (green) and KMO (red)-expressing cells in the spinal cord

Movie 3. NEUN^+^ neurons (green) and KMO (red)-expressing cells in the spinal cord
